# *Pseudomonas aeruginosa* core metabolism exerts a widespread growth-independent control on virulence

**DOI:** 10.1101/850206

**Authors:** Stavria Panayidou, Kaliopi Georgiades, Theodoulakis Christofi, Stella Tamana, Vasilis Promponas, Yiorgos Apidianakis

**Author notes:** Corresponding authors VP and YA. These authors contributed equally to this work (Kaliopi Georgiades, Theodoulakis Christofi, Stella Tamana).

## Abstract

Bacterial virulence may rely on secondary metabolism, but core metabolism genes are assumed to be necessary primarily for bacterial growth. To assess this assumption, we correlated the genome, the transcriptome and the pathogenicity of 30 fully sequenced *Pseudomonas* strains using two *Drosophila* and one mouse infection assay. In accordance with previous studies gene presence-absence does not explain differences in virulence among *P. aeruginosa* strains, but merely between *P. aeruginosa* and other *Pseudomonas* species. Similarly, classical gene expression analysis of highly vs. lowly pathogenic *P. aeruginosa* strains identifies many virulence factors, and only a few metabolism genes related to virulence. Nevertheless, assessing the virulence of 553 core metabolic and 95 random non-metabolic gene mutants of *P. aeruginosa* PA14, we found 16.5% of the core metabolic and 8.5% of the non-metabolic genes to be necessary for full virulence. Strikingly, 11.8% of the core metabolism genes exhibit defects in virulence that cannot be attributed to auxotrophy. The compromised in virulence metabolic gene mutants were mapped in multiple pathways and exhibited further defects in acute virulence phenotypes and in a mouse lung infection model. Functional transcriptomics re-analysis of core metabolism at the pathway level, reveals amino-acid, succinate, citramalate, and chorismate biosynthesis and beta-oxidation as important for full virulence and expression of these pathways indicative of virulence in various strains. Thus, *P. aeruginosa* virulence variation, which to this point remains unpredictably combinatorial at the gene level, can be dissected at the pathway level via combinatorial trancriptome and functional core metabolism analysis.

## Introduction

*Pseudomonas* species have been described as Gram-negative, rod-shaped, facilitative aerobic and polarly flagellated bacteria with some sporulating species ^1^, which are very diverse and able to utilize a variety of organic compounds and colonize a wide range of niches. They can also be pathogenic to humans, animals and plants ^1–3^. *P. aeruginosa* strains stand out as human opportunistic pathogens exhibiting a remarkable capacity to inhabit diverse environments, including soil and water, and the ability to infect multiple model organisms, including insects, plants and mice due to their large genomes and remarkable variety of virulence factors ^4–6^. In humans *P. aeruginosa* causes acute infections in injured, burned and immunocompromised patients and persistent respiratory infections in individuals with cystic fibrosis (CF) ^7^.

Previous comparative analyses of *P. aeruginosa* strain-specific genes revealed that different isolates share >90% of their gene content irrespective of the source of their isolation (soil, wound, urinary tract, lung, blood or eyes) or the type of infection they were found to cause ^3,8,9^. The high virulence of bacterial pathogens is usually attributed to the presence of pathogenicity islands, which are clusters of virulence-related genes often acquired by horizontal gene transfer ^10,11^. The genomic sequence of the highly virulent *P. aeruginosa* strain PA14 revealed a core genome with ∼90% of its genes sharing essentially identical homologues with the *P. aeruginosa* prototypic reference strain PAO1, plus islands of genes related to survival in diverse environments ^12,13^. Despite the striking genomic similarity, *P. aeruginosa* strain-to-strain variation in virulence is vast ^12,14,15^ and cannot be simply explained by the presence of virulence-related genes residing on pathogenicity islands ^3^. This comes against examples of genome decay often described in comparative genomic studies trying to explain the adaptation of pathogens to a host-associated lifestyle ^6,16,17^.

Besides the identification of many virulence factors and some of their immediate regulators, *P. aeruginosa* pathogenicity appears not to follow the same paths for each strain. *P. aeruginosa* strains can evolve, nevertheless, in the lung of cystic fibrosis patients, and some strains, such as CF5, are only weakly pathogenic in models of acute infection ^8,14^. Despite extensive efforts to link gene content with pathogenicity of *P. aeruginosa,* virulence cannot be explained by the presence or absence of single virulence genes ^3,8^. Accordingly, *P. aeruginosa* pathogenicity has been characterized as context dependent, that is, genes required for pathogenicity in one strain may not necessarily contribute to virulence in other strains. We hypothesized that, if not gene presence, then gene expression of virulence factors and their regulators should be able to explain differences in pathogenicity among *Pseudomonas* strains.

In the current study, we compared 30 fully sequenced *Pseudomonas* genomes using large-scale comparisons at the protein sequence level. Primarily we sought to identify differences in the presence and absence of genes that may explain the differences in pathogenicity among these strains. We also established an extensive set of known virulence factors (VFs), the expression rather than the presence of which correlated with highly vs. lowly virulent strains. In addition, to VFs, core metabolism gene expression correlates with pathogenicity, but at the pathway rather than the gene level. At the functional level we estimated the percentage of the core metabolism genes participating in the virulence of *P. aeruginosa* strain PA14 and found it to be at least as high to that of the non-metabolic genes. Despite the known interconnectivity between bacteria core metabolism and growth ^18^, 11.8% of core metabolic genes were contributing to virulence and virulence factor production, while being non-essential for growth in culture or in the host. Our functional transcriptomics approach and analysis at the pathway level reveals the existence of a widespread growth-independent effect of core metabolism modules in virulence.

## Methods

### Bacterial strains

The catalog of the core metabolic genes of *P. aeruginosa* strain PA14 was produced in 2014 using the KEGG Database ^19^. The 553 metabolic gene mutants were picked from the 96-well plates of the publicly available PA14 Transposon Insertion Mutant Library, while the 95 non-metabolic gene mutants were randomly selected from sequential plates of the same library verifying that none of them is linked in the KEGG database with bacterial metabolism (http://ausubellab.mgh.harvard.edu/cgi-bin/pa14/home.cgi). The 30 wild type sequenced *Pseudomonas* strains used in this study are described in **Suppl. Table 1**.

### Infection Assays

#### Intranasal Mouse Lung Infection Assay

The intranasal infection achieves the spreading of the bacteria from the upper airways to the intestine and low airways, thus mimics the pathology seen in acute bacterial pneumonia ^20,21^. PA14 wild type and mutant strains were grown in LB liquid cultures overnight. Cultures were then diluted 1:100 and were grown over day to OD_600nm_: 3.0 (∼3×10^9^ CFU/ml). Bacteria were pelleted and washed twice in sterile saline (0.9 %) and a required dilution was done in order to reach the desired infectious dose of 2×10^7^ CFU/mice. Mice were intranasally infected, under very short and light anesthesia, as previously described ^22,23^, by placing 10μl of a bacterial suspension in each nostril (20μl in total). Mortality counts were taken every day for 7 days.

#### Ethics statement

Animal protocols were approved by the Cyprus Veterinary Service inspectors under the license number CY/EXP/PR.L6/2018 for the Laboratory of Prof. Apidianakis at the University of Cyprus. The veterinary services act under the auspices of the Ministry of Agriculture in Cyprus and the project number is CY.EXP101. These national services abide by the National Law for Animal Welfare of 1994 and 2013 and the Law for Experiments with Animals of 2013 and 2017. All experiments were performed in accordance with these guidelines and regulations.

#### Fly Pricking Assay

Male Oregon R flies were pricked in the thoracic cuticular epithelium with a tungsten needle dipped in a bacteria suspension, as previously described ^24^. The infection mix contained 980 μl ddH_2_O, 10 μl 1M MgSO_4_ and 10 μl of bacteria OD_600_: 3.0. Vials were transferred at 25°C and survival was measured every day. For each mutant we had 2 vials with 20 flies (40 flies in total) for more robust statistical analysis.

#### Fly Feeding Assay

Female Oregon R flies 3-7 days old were starved for 5-6 hours and then were transferred in vials containing the following feeding mix on cotton: 0.5 ml of bacteria OD_600_: 3.0, 1 ml 20% sucrose and 3.5 ml ddH_2_Ο. Vials were transferred at 25°C and survival was measured every day. For each mutant we had 3 vials with 10 flies (30 flies in total) for more robust statistical analysis.

#### Flies survival

For the determination of the mortality rate of the infected flies, daily observation of the experimentations was necessary. Uninfected flies can live up to 60 days; however, infection using *P. aeruginosa* PA14 usually kills flies in less than 10 days. Fly infections were repeated in technical replicates two independent times for each bacterial strain for both assays (oral and wound infection). The fly survival was calculated based on the total number of flies surviving.

#### Strain clustering based on pathogenicity

A survival graph was obtained for the flies infected by each bacterial strain and the survival curves were statistically analyzed using the Kaplan-Meier method ^25^ as implemented in IBM SPSS. The resulting survival curves were compared pairwise in an all-against-all fashion, using a chi square test (χ^2^). Applying the unweighted pair group method with arithmetic mean (UPGMA) agglomerative clustering procedure ^26^ (using the respective χ^2^-values as a pairwise distance) produces a tree-like representation for the strains analysed. This “natural” hierarchical clustering was further used to separate *Pseudomonas* strains in three groups (corresponding to low, medium and high virulence) by cutting the tree in the second bifurcation from the root. Visual inspection of the Kaplan-Meier survival curves and the requirement that the well-characterized avirulent (i.e. CF5) or highly virulent strains (i.e. PA14) should be assigned as “low” and “high”, respectively, led to the designation of labels to the resulting clusters.

### Phylogenetic analysis of *Pseudomonas* strains

We produced a reliable phylogeny of the 30 studied Pseudomonas species/strains which was necessary as a guideline for correlation with the pathogenicity scale. We followed the approach described in Duan *et al.*, 2013 ^27^, the only difference being that *rpoB* and *rpoD* homologs could not be identified in the genome of *P. aeruginosa* 2192. Therefore, a phylogenetic tree was constructed using the sequences of the universally present gyrase B (*gyrB*) and 16S rRNA genes for the 30 studied species. *Escherichia coli* K12 and *Cellvibrio japonicus* strain Ueda107 were used as outgroups. Sequences were retrieved from the Pseudomonas Genome Database ^28^ and the NCBI nucleotide database ^29^ for the outgroups, and were multiply aligned using ClustalO ^30^ with default parameters. The resulting multiple sequence alignments were concatenated using the MEGA software package ^31^ and the data matrix produced was further analysed with MrBayes v3.2 ^32^ (using the default parameters: lset nst=6 rates=invgamma, mcmc ngen=20000 samplefreq=100 printfreq=100 diagnfreq=1000), which performs Bayesian inference of phylogeny. An essentially identical *Pseudomonas* phylogeny tree was built with the complete gene matrix for the 29 species (i.e. excluding *P. aeruginosa* 2192) using the same procedure.

### Comparative genomics versus pathogenicity ranking

#### Sequence analysis and clustering

Protein sequences encoded in the *Pseudomonas* strains of interest were downloaded from the Pseudomonas Genome Database ^33^and were internally codified following the style of the COGENT database for consistency and easy manipulation from computational tools ^34^. Sequences were filtered with CAST ^35^(using default parameters) and converted in database records for the NCBI BLAST suite of tools ^36,37^ as previous studies have shown that filtering sequences in regions of extreme amino acid composition can eliminate most of the false positive results in BLAST searches without sacrificing sensitivity ^38^. In this study, a new, more effective version of the CAST algorithm was used ^39^ offering optimized performance for pan-genome analyses. We used a local installation of BLASTP for performing all-against-all pairwise comparisons (e-value threshold: 10^-6^; Composition Based Statistics: off; masking mode turned: off; all other parameters left to their default values). BLAST results were used for sequence clustering using the MCL software ^40,41^ employed with the default parameters as suggested by the authors for the delineation of *Pseudomonas* protein families.

#### Virulence factor (VF) selection

We collected annotated VFs encoded in the *P. aeruginosa* PAO1 genome based on the relevant section of the Pseudomonas Genome Database. Next, in order to obtain a comprehensive dataset of known VFs for most of the 30 studied species, an exhaustive literature search was manually realized in PubMed using keywords such as, “virulence factor”, “pathogenic capacity”, “secretion system”, in correlation with “*Pseudomonas”* species. Our list was further expanded by consulting the Virulence Factors Database (VFDB) ^42^ as a complementary reference list. In this way, a list containing 254 virulence factors was created and VFs were then separated in five groups according to their function: type II secretion system (19 genes), type III secretion system (37 genes), type VI secretion system (18 genes), quorum sensing (44 genes) and other genes that did not enter any of these categories (136) (**Suppl. Table 2**).

#### Metabolic genes

We identified the proteins annotated to participate in distinct metabolic pathways and categories based on the KEGG database (Kyoto Encyclopedia of Genes and Genomes, www.kegg.jp^19^. For the assessment of RNAseq transcriptomics data (see below) against metabolic pathways, we deliberately used an independent pathway definition provided by BioCyc ^43^, in order to avoid any potential biases due to our initial selection of metabolic genes.

#### Phylogenetic profiles of *Pseudomonas* VFs and metabolic genes

Instead of relying to external resources for inferring homology among proteins encoded in the genomes of interest, we employed the bidirectional-best hit (BBH) approach. Briefly, parsing the all-against-all BLASTP results with custom developed software, we identified cases where a protein *protA* from “Genome A” has *protB* as a best hit (according to the BLASTP score) in “Genome B”, which in turn has *protA* as its best hit in “Genome A”. This procedure rapidly identifies cases of 1-1 orthologs, with the drawback that some subtle cases (e.g. where recent gene duplication has occurred) may be missed ^44^. Moreover, cases of wrongly predicted genes are expected to result in additional proteins that seem to be absent from some genomes of interest. In order to alleviate this drawback, all proteins of interest that appeared to be absent from particular *Pseudomonas* strains after the BBH analysis were further queried using translated BLAST searches (TBLASTN) against the respective target genomes and if reliable hits were found they were recoded as “present”. Phylogenetic profiles for *Pseudomonas* VFs and metabolic proteins were coded in Presence-Absence (PA) tables using a simple +1/-1 scheme (present/absent). PA tables were further visualised using the MeV software ^45^ multiple array viewer feature and were subject to clustering using the k-means method in the R environment ^46^. These clusters were then compared to the groups obtained by the pathogenicity ranking using the Adjusted Rand Index (ARI) ^47,48^ as implemented in the mclust R package. The ARI is a measure for the chance grouping of elements between two clustering’s accompanied by a measure of the statistical significance of the overlap of the groups compared.

### Bacterial-RNAseq analysis and deep sequencing

Bacterial RNA was isolated from LB cultures at OD_600nm_ 1 and 3 in biological replicates using the QIAzol lysis reagent (Qiagen). The quantity of the bacterial RNA samples was measured on a NanoDrop ND1000 system and their quality analyzed on the Agilent 2100 Bioanalyzer system with the Agilent RNA 6000 Nano kit protocol (Agilent Technologies), according to manufacturer’s instructions. Bacterial RNA samples with RNA Integrity Number (RIN) > 7 were chosen for mRNA enrichment. The MICROBExpress™ Bacterial mRNA Enrichment Kit protocol (ThermoFisher Scientific) was performed on 5 – 10μg of bacterial RNA, according to manufacturer’s instructions. The quality and quantity of enriched mRNA was measured using the NanoDrop and the Bioanalyzer systems. After assessment of the enriched bacterial mRNAs, libraries were prepared using the Ion Total RNA-Seq Kit v2 protocol and reagents (ThermoFisher Scientific) according to manufacturer’s instructions.

The enriched mRNAs were fragmented, reverse transcribed and amplified after the addition of a specific (∼10bp) barcode to each sample. The quantity and quality of the prepared libraries was assessed on a Bioanalyzer using the DNA High Sensitivity Kit, according to manufacturer’s instructions. All libraries were diluted to 1nM and subsequently pooled together in pools of eight. 40pM of pooled libraries were further processed for templating and enrichment, and loaded onto the Ion Proton PI™ V2 chips (ThermoFisher Scientific). The procedure was performed on the Ion Chef™ System using the Ion PI™ IC200 Chef kit, protocol and reagents (ThermoFisher Scientific), according to the manufacturer’s instructions. Sequencing of the loaded chips was performed using the Ion PI™ Sequencing 200 V3 kit on an Ion Proton™ system (ThermoFisher Scientific), according to the manufacturer’s instructions. Initial analysis, sample/barcode assignment, and genome mapping was performed on the Ion Proton server.

Initial attempts to map reads on the genomes of origin gave poor results with respect to the number of unassigned RNAseq reads (not shown). Instead, we mapped all reads to the annotated *P. aeruginosa* PA14 genome as a reference (GCF_000014625.1_ASM1462v1_genomic.gff). Reads per gene were counted using featureCounts ^49^ and those corresponding to rRNA genes were in silico depleted to avoid skewing the statistics of independent samples prior to processing with DESeq2 ^50^ for the detection of differentially expressed genes.

#### Pathway enrichment analysis

Relevant gene lists were uploaded to BioCyc SmartTables. Pathway enrichment was performed using the respective BioCyc functionality using the Fischer exact test and the Parent-Child Intersection method that has been shown to be robust against false positives ^51^. Multiple testing correction was performed with the Benjamini-Hochberg procedure. Reported adjusted p-values were considered significant when p_adj_<0.05.

#### Cluster analysis of patterns of differential gene expression

Cluster analysis and visualization of patterns of up-/down-regulation was performed with ClustViz ^52^ on a matrix where up-/down-regulation of each gene was represented with +1/-1, respectively. Zero values were used to denote the absence of significant differential expression.

### Minimal Media Assays

Bacteria that were grown in 3 ml LB O/N cultures, 500 μl were centrifuged at 8000 rpm for 2 min, supernatant was removed, and the pellet was diluted in minimal media. From this mix, 1:100 was left to grow in minimal media on a culture rotator at 37°C. For each mutant we had duplicates and three time points of the optical density OD_600_ were taken. Minimal medium solution of 50 ml consisted of 10 ml 5xM9, 100 μl 1 Μ MgSO_4_, 1 ml 20% glucose and 39 ml ddH_2_O. Minimal medium with 5% fly extract was produced by adding at the above solution extract from grinded flies to a 5% final concentration.

### Fly Colonization Assays

#### Wound Colonization Assay in flies

The colonization ability of the pricking assay-selected metabolism mutants was examined by counting CFUs from grinded flies previously injected with an amount of 10^2^ bacteria. For each mutant and for the wild type strain, 20-25 flies were injected dorsoventrally as previously described ^24^ and then transferred into vials with fresh food (at 25°C). The next day (18-24 hours after), plates were cultured with a solution of grinded flies to count CFUs and observe the colonization ability of each mutant strain, compared to the wild type strain.

#### Intestinal Colonization Assay in flies

The colonization ability of the feeding assay-selected metabolism mutants was examined, using Oregon R flies 3-7 days old, previously starved for 5-6 hours and then fed with bacteria OD_600_: 3.0 for one day. The flies were next transferred into 50 ml eppendorfs with 12 holes (1.2 mm in diameter) on the lid. Flies were able to reach food (Whatman filter paper disc soaked with 200 μl of 4% sucrose and 10% LB set on the lid and covered with parafilm) only by the holes of the lid. Flies were transferred in clean 50 ml eppendorfs every day for 3 days to avoid contamination. The third day, plates were cultured with a solution of grinded flies to count CFUs and observe the colonization ability of each mutant strain, compared to the wild type strain.

### Quantitative reverse transcription real-time PCR

RNA isolation was performed from log-phase cells grown to an OD_600_ of 1.0 or 2.0 using the QIAzol Lysis Reagent. Briefly, 500 μl of bacteria OD_600_ of 1.0 or 2.0 (∼10^9^) were spun down at 8000 rpm for 3 minutes. The pellet was dissolved by adding 500 μl of QIAzol and by pipetting up and down at 60°C for 10 minutes. Next, 100 μl CHCl_3_ was added and tubes were inverted for 15 seconds. After a 5 minutes incubation at room temperature, the supernatant (300 μl) was selected in a new tube, by a full speed centrifugation at 4°C for 15 minutes and then mixed with 300 μl iso-propanol by inverting the tubes. Tubes were then let on bench for 5 minutes and after a step of a full speed centrifugation at 4°C for 10 minutes, the pellets were washed out with 500 μl of 70% EtOH. Finally, after a centrifugation at 4°C for 3 minutes and air drying, pellets resuspended in 20-50 μl RNase free H_2_O by pipetting and stored at −80°C. The bacterial RNA was reverse transcribed into complementary DNA (cDNA) using the PrimeScript RT reagent Kit (Perfect Real Time) (Takara: RR037A) according to the manufacturer’s instructions. Real-time PCR was performed, using a Bio-Rad CFX1000 thermal cycler and Bio-Rad CFX96 real-time imager with primer pairs listed below and iQ SYBR green supermix (Bio-Rad).

#### Target gene primers sequence (5’-3’)

“F” and “R” indicate the direction (forward or reverse) of each primer

**rplU** F: ATGGCGAAGACGTGAAAATC, R: GAACTTGATGATGCGGACCT

**clpX** F: GTTCGGTCTTATCCCCGAGT, R: AACAGCTTGGCGTACTGCTT

**exsC** F: CACCGTTTCGATCTGCATTTCG, R: CGAGAATCTGCGCATACAACTG

**exoT** F: GCCGAGATCAAGCAGATGAT, R: TTCGCCAGTCTCTCCTCTGT

**pa1L** F: GGTTGCACCCAATAATGTCC, R: CCAATATTGACGCTGAACGA

**pilA** F: CAGAGGCGACTGGTGAAATC, R: AGGGTAGAGTCAGCCGGAAT

Results are from two independent experiments performed in biological triplicates. All samples were normalized to the expression of the housekeeping genes *rplU* and *clpX* via the Pfaffl method ^53^.

### Motility Assays

#### Swarming motility

Swarming motility was assessed using petri dishes each containing exactly 20 ml of medium with 5 g/l Bacto-agar (Difco), 8 g/l Nutrient Broth (Difco) and 5 g/l Dextrose. Bacterial cultures with the strains of interest were grown overnight in LB medium and 2 μl from each culture was added at the center of a swarming plate. The plates stayed open until the droplet was fully absorbed by the agar and then incubated for 24 hours at 37°C. The diameter of the swarming zone on the plate was measured, and photos were taken.

#### Twitching motility

Twitching motility was assessed using petri dishes containing 1.0% Bacto agar and 20 g/L LB broth. After the agar was solidified, the indicative strains were stabbed at the bottom of the plates with a sterile toothpick. The plates were incubated at 37°C for 48 hours. The ability of the bacteria to adhere and form biofilms was examined by removing the agar, washing the unattached cells with water and staining the attached cells with crystal violet (1%). The stain solution was removed by carefully washing the plates with water. The diameter of the twitching zone on the plate was measured, and photos were taken.

## Results

### Pathogenicity-based grouping of *Pseudomonas* strains

To correlate the pathogenicity of 30 fully sequenced *Pseudomonas* strains (18 *P. aeruginosa* and 12 other *Pseudomonas* strains; **Suppl. Table 1**), we examined their virulence in *Drosophila melanogaster*, using an oral and a wound infection assay that impose two distinct types of acute infection (**Figure 1 A, B**) ^24,54^. Kaplan-Meier survival analysis with a log rank test and pairwise comparison over strata, was used to analyze the results of fly survival after the infection. Even though it is usually intuitive to group bacterial strains based on visual inspection of their survival curves, this procedure is subjective. To objectively partition the strains under study based on their pathogenicity, the 30 bacterial strains were classified in three groups (low, medium and high), depending on the severity of their virulence phenotype (**Figure 1 A, B** and **Suppl. Table 1**), using hierarchical clustering (see Materials and Methods; **Suppl. Tables 3-4**). We used the PA14 and CF5 strains to define the “high” and “low” virulence categories, respectively, in both assays, in accordance with published data ^8,14^.

**Figure 1.**
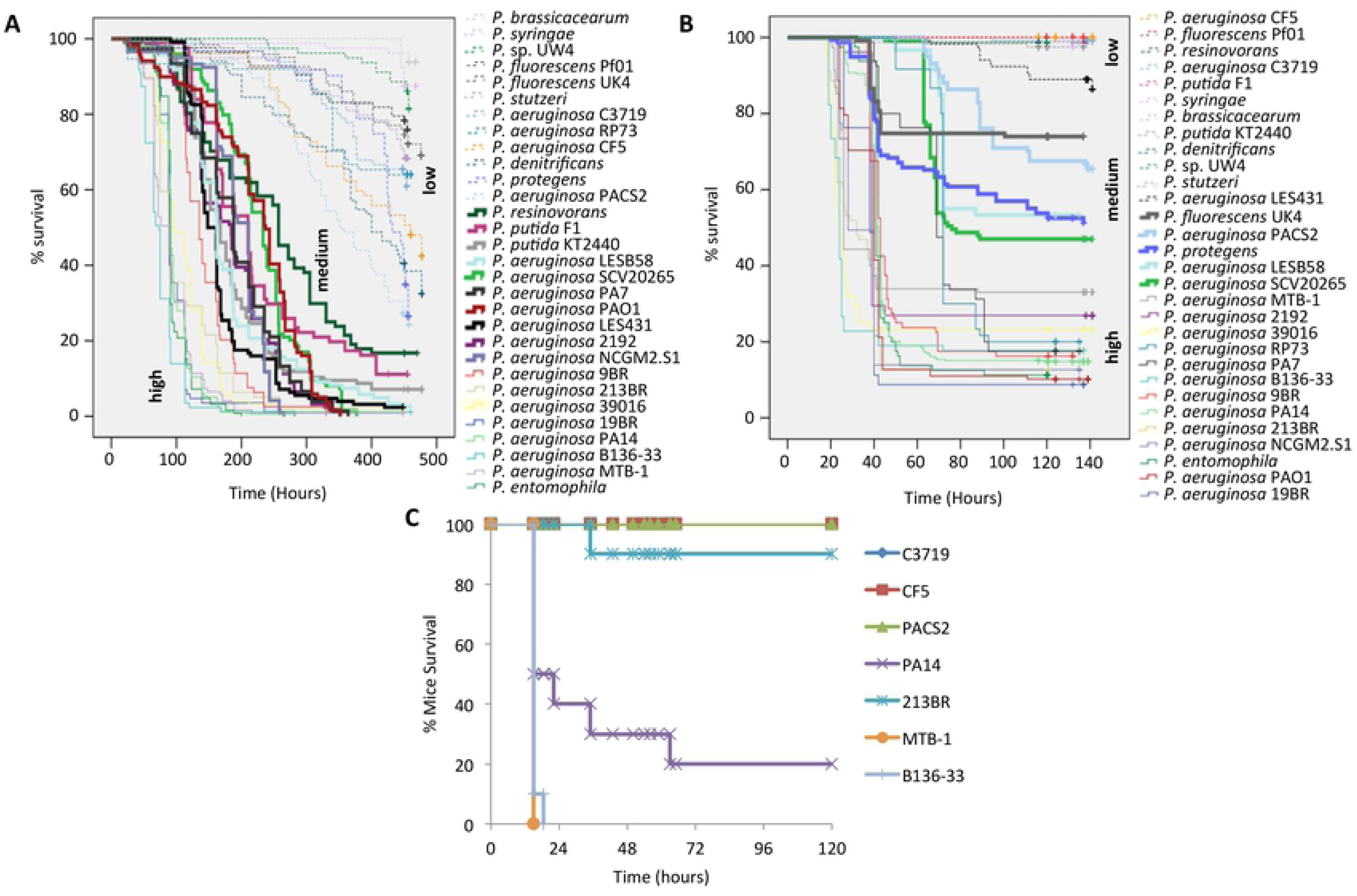
Comparative survival of all 30 *Pseudomonas* strains upon oral and wound infection in flies and lung infection in mice. **(A, B)** Three groups are distinguished, each representing species that are either high, medium or low virulent in fly oral (A) and wound (B) infection. Thin lines represent high virulent strains, thick lines represent medium virulent strains and dashed lines represent low virulent strains. Time is measured in hours. (**C**) Mouse survival (%) after intranasal mice infection with 3 lowly and 4 highly virulent *P. aeruginosa* strains. Twenty-microliter aliquots of bacterial solution containing 2×10^7^ bacteria was administered intranasally to each mouse. Infected mice were monitored for 5 days (n=10).

Non-*P. aeruginosa* strains tend to consistently group in the “low” virulence cluster, with a notable exception of *P. entomophila* presence in the “high” virulence cluster. *P. entomophila* is a known entomopathogenic bacterium, able to infect and kill insects, including *Drosophila* ^55^. *P. aeruginosa* strains B136-33, MTB-1, PA14, 213BR, 19BR, 9BR and 39016, on the other hand, were consistently virulent, while strains CF5 and C3719 consistently low in virulence regardless of the infection assay. Some *P. aeruginosa* strains were less consistently grouped between assays, such as PAO1, which was grouped with the “high” and “medium” virulence cluster in the wound and oral infection assays respectively. To more rigorously select highly and lowly virulent strains we used a model of acute intranasal mouse lung infection examining the mortality rate of four of the highly (B136-33, MTB-1, PA14, 213BR) and the three most lowly (C3719, CF5, PACS2) in *Drosophila* virulence *P. aeruginosa* strains. We found that all but one (213BR) of the 7 tested strains retained their virulence potential in the acute mouse lung infection model (**Figure 1** **C**).

### Presence-Absence of single genes cannot explain *Pseudomonas* pathogenicity

We inferred a phylogenetic tree of all the 30 *Pseudomonas* strains based on the concatenated alignments of two core housekeeping genes, the 16S rRNA and *gyrB* (**Suppl. Figure 1**), which was in agreement with the trees constructed using four housekeeping genes or 1679 orthologs per Duan *et al.*, 2013 ^27^. Our results were also in agreement with previous observations that place *Pseudomonas* sp. UW4 within the *P. fluorescens* clade, despite the fact that phylogenetic analysis of 16S rRNA sequences alone indicates a closer relationship to the *P. putida* clade.

Next, we created a database of virulence factors (VFs) containing 254 sequences (see Methods). We classified them according to their functions in five groups: type II secretion system (T2SS; 19 genes), type III secretion system (T3SS; 37 genes), type VI secretion system (18 genes), quorum sensing (44 genes) and other (136) (**Suppl. Table 2**). For each category a BBH BLASTP search allowed us to detect the phylogenetic profiles of VFs across the *Pseudomonas* species. Five multiple array viewers (one for each functional category) were obtained indicating presence-absence of VFs. A sixth multiple array viewer was obtained by merging all VFs together (**Figure 2**). All VFs found to be absent in a particular genome were manually re-examined with a TBLASTN analysis to exclude cases where the observed absence was the result of erroneous gene-finding or incomplete annotation. We find that most VFs are present in all *P. aeruginosa* strains, even in the less virulent strains, including stain CF5 (**Figure 2** and **Suppl. Table 1**). Most *P. aeruginosa* VFs are absent in non-*P. aeruginosa* strains (**Figure 2** and **Suppl. Table 1**). It is possible though that other (non-homologous) genes/proteins compensate for the same functions in non-*P. aeruginosa* strains.

**Figure 2.**
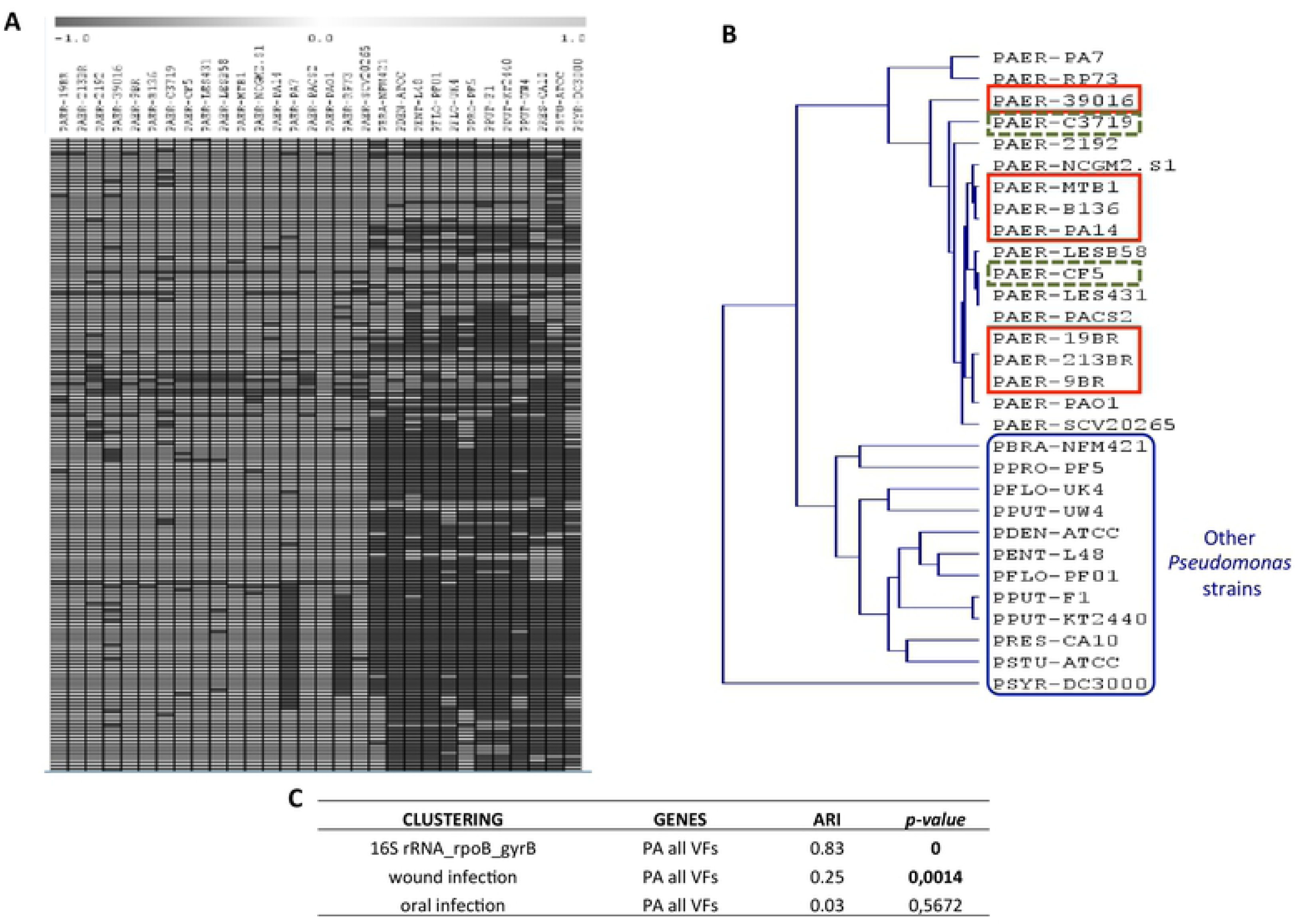
Virulence factor Presence-Absence follows phylogeny rather than pathogenicity. (**A**) Part of a multiple array viewer based on presence/absence of all virulence factors. Present genes are shown in light grey, absent genes are shown in dark grey. (**B**) Clustering based on presence/absence of all virulence factors. The 2 lowly virulent in both fly infection assays *P. aeruginosa* strains are shown in dashed-line boxes; and the 7 high virulent in both assays *P. aeruginosa* strains in full-line boxes. (**C**) Phylogeny rather than pathogenicity of the 30 Pseudomonas spp. correlates best with Presence-Absence (PA) of virulence factors (VFs).

For type II secretion system (T2SS), all *P. aeruginosa* strains have all genes except from strain PA7 missing the toxA gene, as was previously demonstrated ^12^. However, non-*P. aeruginosa* strains are missing almost all T2SS factors. For example, *P. putida* KT2440 is known for lacking the toxA gene, while the xcp family is replaced by the Gsp family in this strain ^56^.

For type III secretion system (T3SS), four *P. aeruginosa* effector proteins have been functionally identified so far, exoU, exoS, exoT and exoY ^57^. Surprisingly, most *P. aeruginosa* strains possess either the *exoS* or the *exoU* gene but not both, with the exception of strain PA7 that has none. The strain PA7 has no T3SS at all and a compromised virulence phenotype has been attributed to this strain ^12^. The *P. aeruginosa* 39016 strain is missing five T3SS genes (excE, excC, pcr3, popD, pscE), the LESB58 is missing three genes (pcrH, excE, pscP) and the CF5 strain, known as a lowly virulent *P. aeruginosa* strain is only missing one gene (pscQ). One T3SS gene is also missing from strain 2192 (excC) and strain SCV20265 (pscE). Just like for T2SS, T3SS is nearly entirely missing from all non-*P. aeruginosa* strains, e.g. *P. putida* was known for not having a T3SS ^56^.

Regarding the type VI secretion system, the 39016 strain is missing three genes (hcpC, tagT, vgrG1) and the C3719 strain is missing only one gene (dotU). Interestingly, *P. fluorescens* has the T6SS gene hcp and *P. protegens* has the vgrG, hcp, lip, icmF, dotU, clpV, ppka and pppA genes. Moreover, the *P. aeruginosa* quorum sensing genes pzA2, phzB2, rhlI and rhlR involved in phenazine biosynthesis are missing, not only from the non-*P. aeruginosa* strains, but also from various *P. aeruginosa* strains, including the low in virulence srtrains CF5 and C3719 (**Suppl. Table 2**).

We proceed by clustering *Pseudomonas* strains based on the presence-absence of specific VFs for the five VF functional classes and for the complement of VFs. The resulting clusters were further analysed using the Adjusted Rand Index (see Materials and Methods) to reveal hidden correlations to (a) overall *Pseudomonas* phylogeny, (b) VF gene content, (c) metabolic pathway gene content, (d) pathogenicity ranking according to the wound infection assay, and (e) pathogenicity ranking according to the oral infection assay. The results showed that both primarily phylogeny, but also pathogenicity to wound infection, correlate with VF gene content, indicating that *Pseudomonas* phylogeny rather than pathogenicity of the 30 *Pseudomonas* spp. should account for VF gene content differences. Moreover, the phylogeny of the 30 strains correlates well with: (i) their virulence classification in the wound infection, but not in the oral infection assay; (ii) VF gene content; and (iii) major metabolic pathway gene content. To eliminate the impact of the presence of the phylogenetically more distant non-*P. aeruginosa* strains in the results, the analysis was repeated with only the *P. aeruginosa* strains and no correlation was evident for any comparison. Thus, we reconfirm earlier works mentioning that the difference in virulence potential within the genus *Pseudomonas* cannot be attributed to the presence-absence of specific genes or gene functional groups ^8^, but rather to broader interspecies differences yet to be determined.

### Gene expression of selected VFs, but not of core metabolism genes, correlates with pathogenicity

To identify differentially expressed genes relating to pathogenicity we performed a transcriptome analysis of the 6 *P. aeruginosa* strains validated in mice as “high” (B136-33, MTB-1, PA14) or “low” (C3719, CF5, PACS2) in virulence. RNA was extracted from bacteria grown at mid exponential and early stationary phase (optical density OD_600nm_ 1 and 3, respectively). We selected genes differentially regulated in the same direction (either over- or under-expressed) between all pairs of “high” vs. “low” in virulence strains. Nine of them were differentially expressed at both growth phases (8 of which were overexpressed in highly virulent strains). At OD_600nm_ 1 fifteen genes were overexpressed in all highly pathogenic strains and 6 were overexpressed in all the lowly pathogenic strains. At OD_600_ 3 the highly and the lowly pathogenic strains showed overexpression for 52 and 2 genes, respectively (**Figure 3 A**). Overall 20 VFs (most of them being quorum sensing and T3SS related), 10 metabolism genes, 5 transcriptional regulators and a number of hypothetical proteins were found differentially expressed (**Suppl. Table 5**).

**Figure 3.**
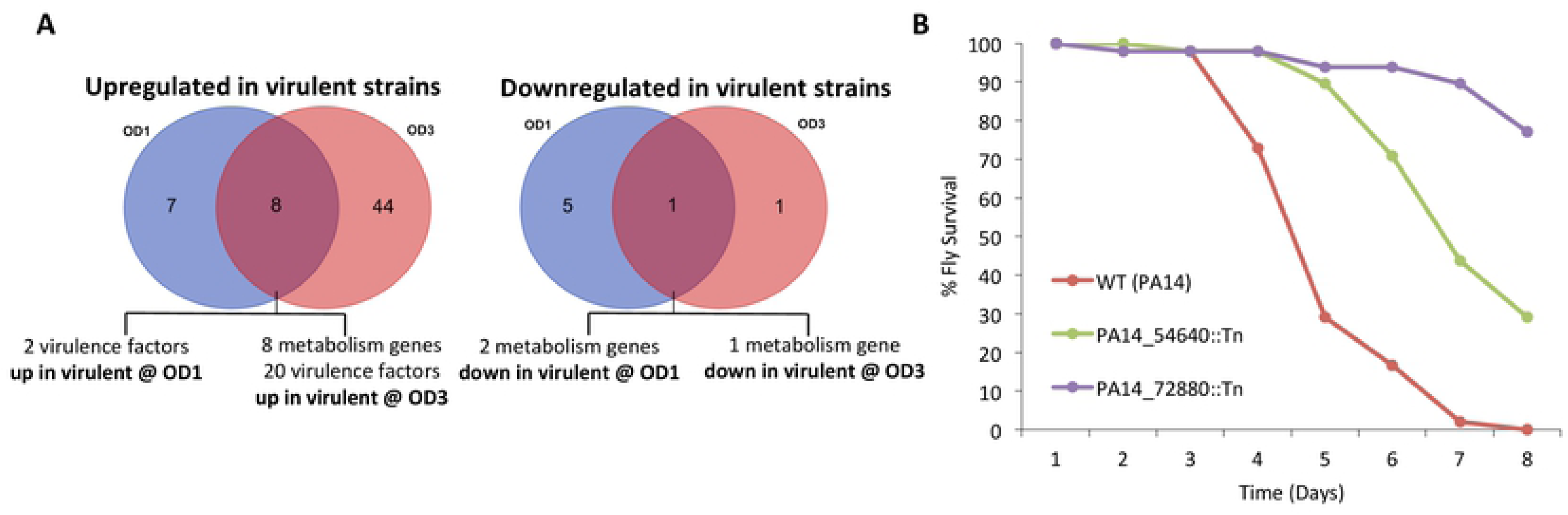
Genes differentially expressed between 3 lowly and 3 highly virulent *P. aeruginosa* strains. (**A**) VENN diagram of the 66 differentially expressed genes identified by RNAseq transcriptomics analysis using http://bioinformatics.psb.ugent.be/webtools/Venn/. At OD1, 15 genes were up (in blue) and 6 genes were down (in yellow) in all 3 highly vs. all 3 lowly pathogenic species. At OD3, 52 were up (in green) and 2 genes were down (in red) in all 3 highly vs. all 3 lowly pathogenic species. (**B**) Mutants for all 10 general metabolism differentially expressed genes were tested for virulence in the fly oral infection model, and 2 of them were found attenuated.

Among the virulence genes overexpressed in highly pathogenic species, ExoU and SpcU were found overexpressed in the highly pathogenic species. This is because they do not exist in the lowly pathogenic species assessed. Furthermore, out of the 18 *P. aeruginosa* species, 5 have ExoU (PA14, B136-33, MTB-1, NCGM2.S1, 39016) and all of them have next to them SpcU, a chaperone required for efficient secretion of the ExoU cytotoxin ^58^, with the exception of 39016 that has no SpcU. Hypothetical proteins overexpressed in highly pathogenic species include: PA14_20600, a DUF3313 domain-containing protein (PFAM ID: PF11769), a domain defining a family of putative bacterial lipoproteins; PA14_04710, a 4Fe-4S ferredoxin that, like other ferredoxins, mediates electron transfer in a wide variety of metabolic reactions ^59^; PA14_30690, a fimbrial protein playing a role in cell adhesion; and PA14_01120 that maps to the antitoxin Tsi6 of the structurally characterized Tse6::Tsi6 complex, a toxin-antitoxin system and is related to T6SSs ^60^. Quorum sensing genes, such as, the phenazine biosynthesis genes *phzB1*, *phzC1*, *phzE1*, *phzM* and *phzS*, were consistently overexpressed among all the highly virulent strains. Thus, contrary to the inability of VF presence-absence profiles to distinguish highly from lowly pathogenic *P. aeruginosa* strains, our data indicate that overexpression of particular VFs is indicative of virulence potential. However, VF regulation is key to our understanding of bacterial virulence, therefore, additional factors need to be investigated.

Among the 10 metabolic genes, 8 were found upregulated in the highly virulent strains, but mutations in none of them exhibited defects in virulence in flies. Mutations, on the other hand, in the 2 downregulated in highly virulent strains metabolic genes exhibit clear defects in virulence (**Figure 3 B**). Of the latter, PA14_54640 (*dspI*) is a putative enoyl-CoA hydratase involved in biofilm dispersion. Its mutation produces high biofilm formation, a condition that may promote chronicity to the expense of acute virulence ^61,62^. The second downregulated in highly virulent strains metabolic gene PA14_72880 is the putative short chain dehydrogenase involved in fatty acid biosynthesis. Mutations of its orthologs in PAO1 reduce biofilm formation, but also swarming motility and LasB protease activity ^63^. The fact that a number of metabolic genes appear among the consistently differentially expressed genes (Fig. 3) begs the question on the broader contribution of metabolism in *P. aeruginosa* virulence ^18^.

### Relative contribution of core metabolic and non-metabolic *P. aeruginosa* PA14 genes to virulence

To functionally assess the contribution of core metabolism genes of *P. aeruginosa* to virulence, we took advantage of the PA14 unigene Transposon Insertion Mutant Library. The PA14 strain is highly pathogenic and well annotated. Its genome contains 6,537,648 nucleotides and 5,977 annotated genes ^3,19^ 794 of which were annotated as metabolic in the KEGG database at the early phase of this project in 2014 (961 as of October 2019). We investigated the virulence in flies of 553 core metabolism gene mutants from the PA14 mutant library corresponding to 482 core metabolism genes and 95 randomly selected mutants corresponding to 94 non-metabolic genes in the pricking (wound) and feeding (oral) infection fly assays. To avoid false negatives, we manually screened 3 replicates of 10 wild type orally infected and 2 replicates of 20 wild type wound infected flies with each of the 648 PA14 mutant strains. The time of 50% fly death (LT50%) was measured for each mutant in each assay and a Z-score analysis was used to select those with a standard deviation >1 i.e. those mutants allowing more fly survival upon infection (**Figure 4 A-D**). To avoid false positives, selected mutants were retested for virulence and the number of the attenuated mutants was finalized after performing Kaplan-Meier survival analysis with a log rank test. Considering both assays, 16.2% (78/482) of the metabolic and 8.5% (8/94) of the non-metabolic *P. aeruginosa* genes, were found virulence-defective in flies (**Figure 5 A**). Even though the number of strains tested only marginally fails to statistically support that more metabolic than non-metabolic genes are implicated in virulence (Fisher exact test: p=0.0583), the numbers clearly suggest that metabolism has a considerable contribution in *P. aeruginosa* virulence.

**Figure 4.**
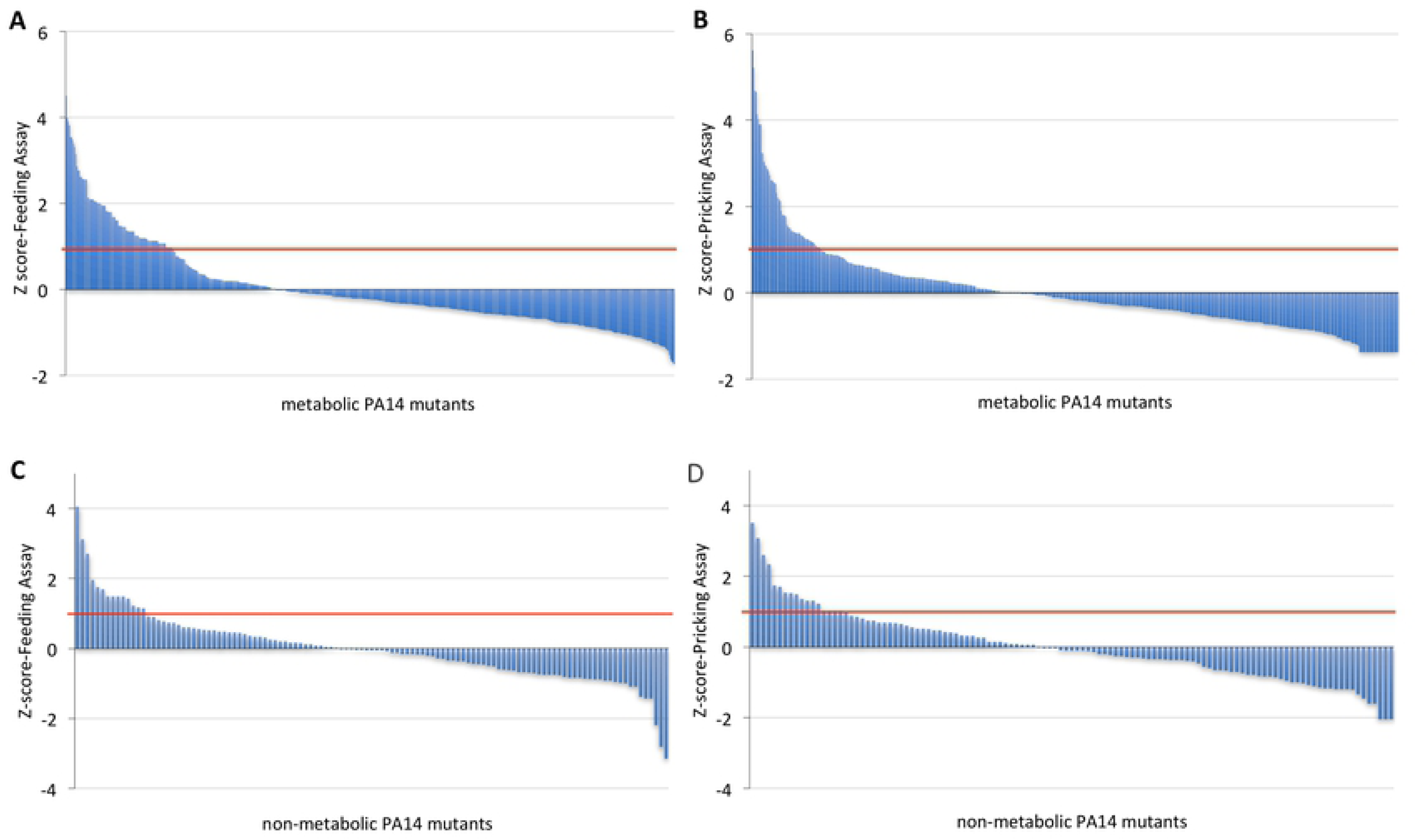
Screen results for 482 metabolic and 94 non-metabolic PA14 genes using two Drosophila infection assays (feeding and pricking). (A-B) Z-score analysis of the fly survival after infection of flies with 553 PA14 metabolic mutants (corresponding to 482 genes) using two independent assays: Feeding Assay **(A)** and Pricking Assay **(B)**. The time of 50% fly death (LT50%) was assessed for each mutant and condition and a Z-score analysis was used to select those with a score >1 for any of the two assays, from which upon retest, 78 were found significantly attenuated in virulence per Kaplan-Meier survival analysis with a log rank test. **(C-D)** Z-score analysis of the fly survival after infection of flies with 95 randomly selected non-metabolic PA14 mutants (corresponding to 94 genes) using two independent assays: Feeding Assay (C), Pricking Assay (D). The time of 50% fly death (LT50%) was assessed for each mutant and condition and a Z-score analysis was used to select those with a score >1 for any of the two assays, from which upon retest, 8 were found significantly attenuated in virulence per Kaplan-Meier survival analysis with a log rank test.

**Figure 5.**
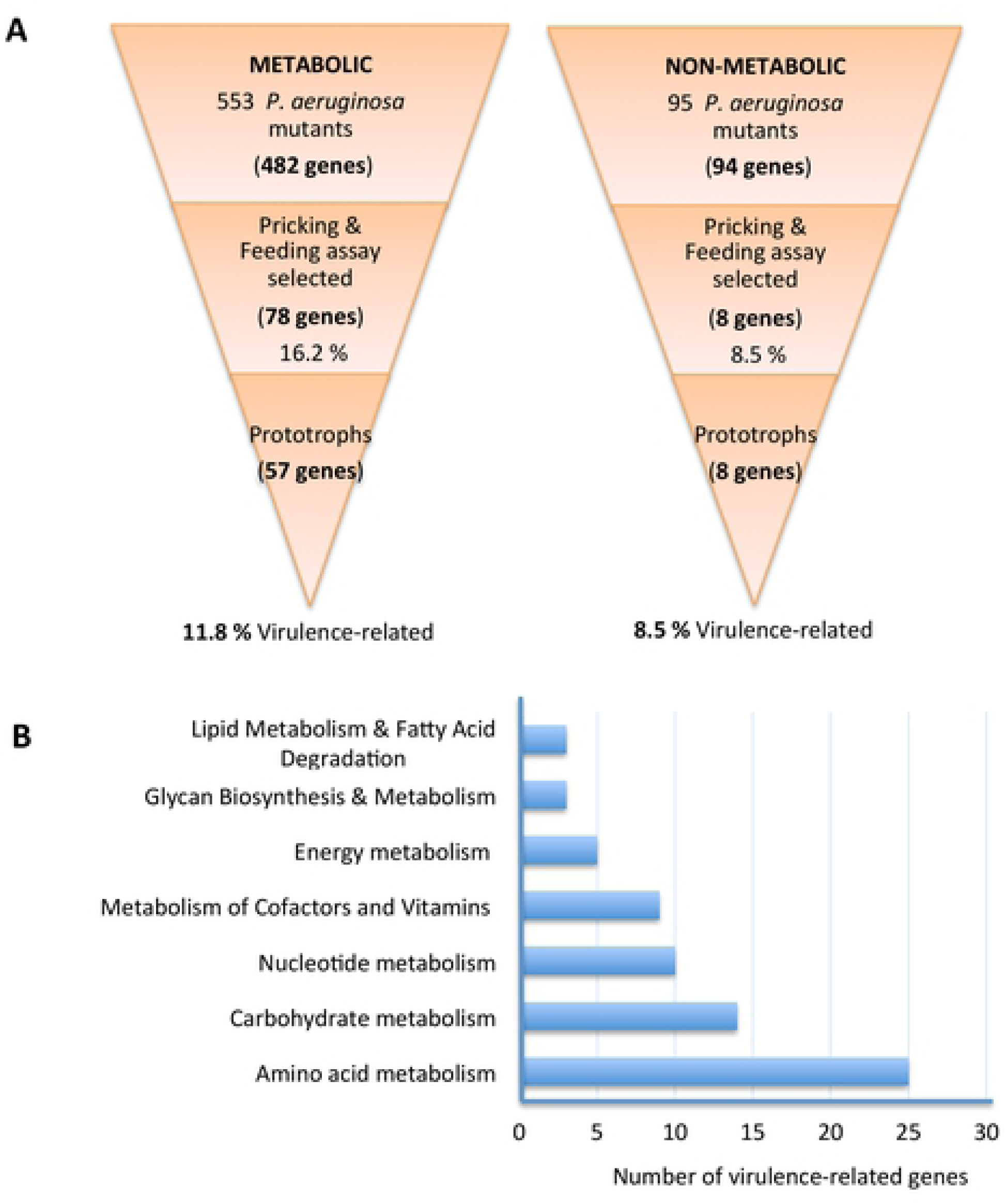
(A) *P. aeruginosa* core metabolism genes are abundantly represented among genes required specifically for full virulence. 553 *P. aeruginosa* metabolic Tn-mutants corresponding to 482 core metabolism genes and of 95 randomly selected non-metabolic *P. aeruginosa* Tn-mutants corresponding to 94 non-metabolic genes were screened in the wound and oral infection *Drosophila* assays. 16.2 % of the core metabolism and 8.5 % of the non-metabolic *P. aeruginosa* genes were selected as required for full virulence. Examination of growth capacity of the selected mutants in minimal media and in the host revealed the percentage of core metabolism genes dispensable for growth, i.e. important specifically for full virulence, is at least as high as that of the non-metabolic genes. (**B) Assignment of the 57 PA14 virulence-related metabolic genes in metabolic categories.** The 57 core metabolism *P. aeruginosa* gene mutants that grow normally in minimal media and/or in the host belong in or more of 7 general core metabolism categories. Each category is represented by at least 3 virulence-related genes, while nucleotide, carbohydrate and amino acid metabolism categories by 10 or more.

### Growth assessment of the virulence-defective PA14 metabolic mutants

The identified metabolism gene mutants might be impaired in growth (exhibit auxotrophy) rather than being directly involved in virulence factor production. Therefore, we sought to determine which of the 78 attenuated in virulence PA14 metabolic gene mutants are growth-essential in culture or necessary for host colonization during infection. We assessed growth in two types of minimal media (without or with fly extract as an extra nutrient source), as well as, the level of colonization during infection at the respective infection model (wound or oral) in which each of these mutants was found attenuated in virulence.

Using a standard minimal medium we initially identified 34 out of the 78 core metabolic mutants as prototrophs for being able to grow similarly to the wild type strain (**Table 1**). The remaining 44 core metabolic mutants did not grow at all in this medium, requiring additional nutrients for their growth. Nevertheless, if the needed nutrients are available in flies during infection, the lack of virulence should be due to a direct effect of the mutants in virulence rather than in growth. To assess this possibility, we inoculated minimal media supplemented with 5% fly extract with each of the 44 core metabolic mutants. This medium offers all the nutrients the bacterium can find in flies in the absence of the host defense against the bacteria. We found 12 of the 44 PA14 metabolic mutants to grow similarly to the wild type and considered them conditional prototrophs (**Table 2 - Group A, B**). All 8 virulence-related non-metabolic mutant strains were also able to grow in this medium (**Suppl. Figure 2 A, B**) and we did not test them any further.

**Table 1.** Thirty-four virulence-related prototrophs based on growth in glucose minimal media. From the 78 virulence-defective PA14 metabolic mutants, 34 can grow like the wild type in the M9 medium, indicating a connection of the corresponding genes with virulence.

|  | Metabolic PA14 mutants | *In vitro* growth | *In vivo* growth (colonization) |  |
| --- | --- | --- | --- | --- |
| # |  | Minimal medium (M9) | Intestinal | Wound |
| 1* | PA14_00120<br>(lipid A biosynthesis lauroyl acyltransferase) | + | - | - |
| 2 | PA14_04320 (ilvA1) | + | + | NT |
| 3 | PA14_07600 (folk) | + | + | NT |
| 4* | PA14_11250<br>(hypothetical protein) | + | - | - |
| 5 | PA14_11810<br>(aldehyde dehydrogenase) | + | + | NT |
| 6* | PA14_14680 (suhB) | + | - | - |
| 7 | PA14_17450 (surE) | + | + | NT |
| 8* | PA14_22050 (htrB) | + | - | - |
| 9 | PA14_20670<br>(glutamine synthetase) | + | + | NT |
| 10 | PA14_29860 (nuoM) | + | + | NT |
| 11 | PA14_29980 (nuoE) | + | + | NT |
| 12 | PA14_29990 (nuoD) | + | + | NT |
| 13 | PA14_31580<br>(acyl-CoA dehydrogenase) | + | + | NT |
| 14 | PA14_32690 (gtdA) | + | + | NT |
| 15 | PA14_38510 (hmgA) | + | + | NT |
| 16* | PA14_40980<br>(enoyl-CoA hydratase) | + | + | + |
| 17 | PA14_44070 (gltA) | + | + | NT |
| 18 | PA14_45010 (hyi) | + | + | NT |
| 19 | PA14_52050 (purN) | + | - | NT |
| 20 | PA14_52610<br>(hypothetical protein) | + | + | NT |
| 21 | PA14_52800 (acsA) | + | + | NT |
| 22 | PA14_65320 (miaA) | + | + | NT |
| 23 | PA14_66670 (ponA) | + | + | NT |
| 24 | PA14_68580 (pckA) | + | + | NT |
| 25 | PA14_70160 (bioA) | + | + | NT |
| 26 | PA14_71970 (wbpW) | + | + | NT |
| 27 | PA14_72850<br>(glutamine synthetase) | + | + | NT |
| 28 | PA14_00020 (dnaN) | + | NT | + |
| 29 | PA14_07170 (epd) | + | NT | + |
| 30 | PA14_08780 (rpoC) | + | NT | + |
| 31 | PA14_31700 (pgsA) | + | NT | + |
| 32 | PA14_32670<br>(hypothetical protein) | + | NT | + |
| 33 | PA14_41020 (adi) | + | NT | + |
| 34 | PA14_63990 (speA) | + | NT | + |
| Feeding assay-selected: 1-27; Those selected in both assays are shown with asterisk (*);<br>Pricking assay-selected only: 28-34; NT: Not Tested |  |  |  |  |

**Table 2.**
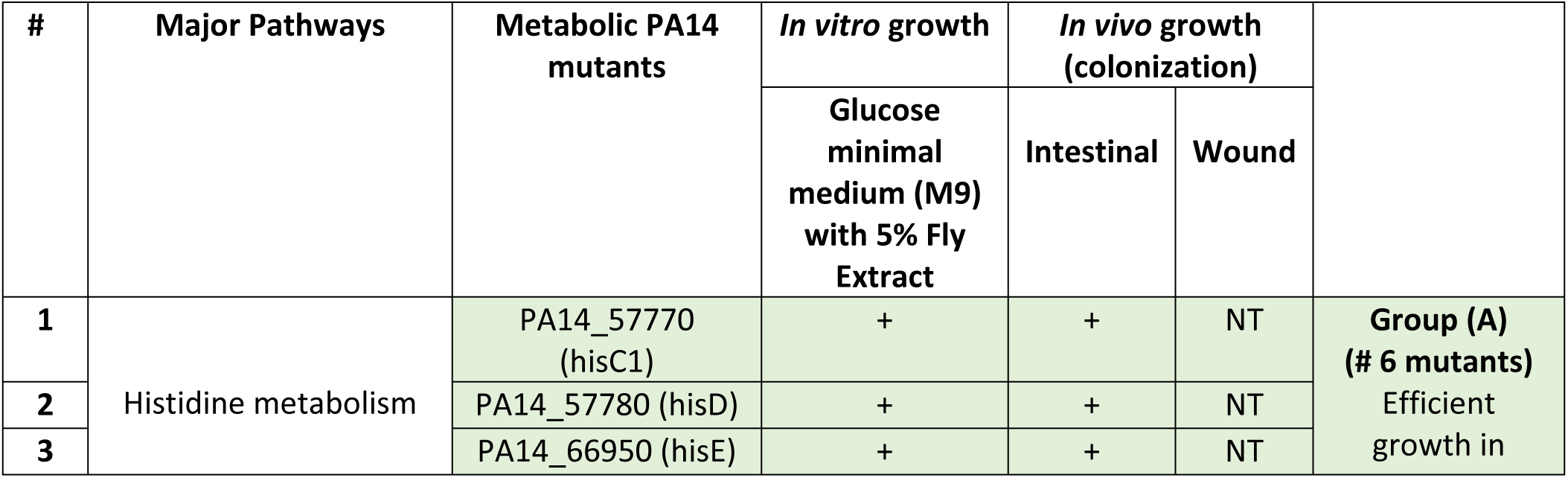
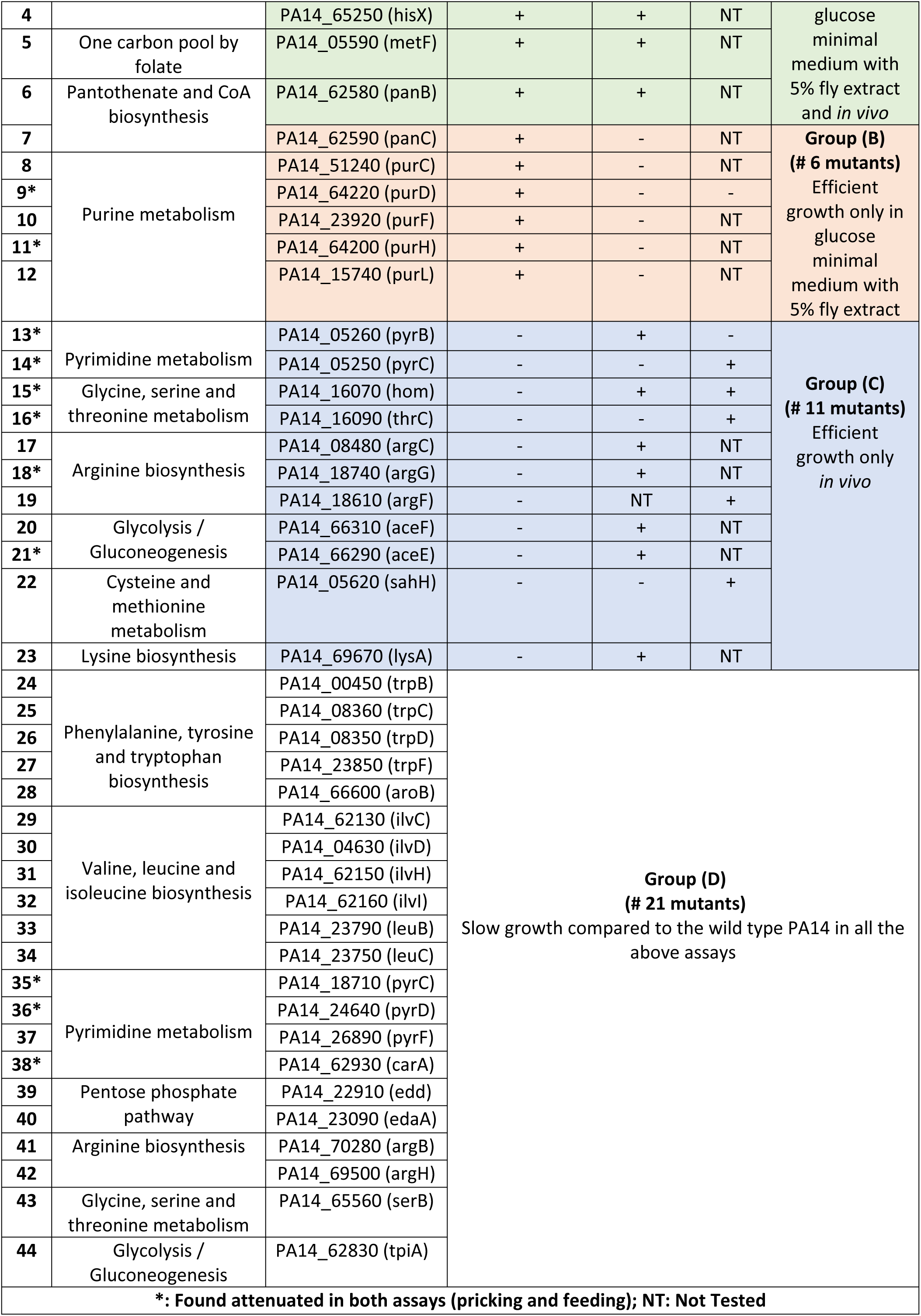
Forty-four virulence-defective mutants: 23 conditional prototrophs & 21 auxotrophs. These metabolic mutants are unable to grow in glucose minimal medium (M9) but 23 of them can conditionally grow *in vitro* and/or *in vivo*, like the wild type strain, if they find in the medium or in the host the missing compounds needed for their growth. The rest 21 do not grow like the wild type in any tested condition, probably because the missing compounds needed for their growth, are not available in the conditions we used.

Moreover, we sought to examine the 78 metabolic mutant strains during initial colonization, to identify auxotrophic strains able to colonize the flies, but also to study the colonization ability of the prototrophs. We adapted our wound and intestinal infection assays to assess colonization efficiency rather than fly survival. For the intestinal colonization assay we infected flies for only one day with all flies being subsequently transferred to tubes with clean food, every day for 3 days. For the wound colonization assay we injected the bacteria in the flies instead of pricking them with a tungsten needle to bypass the fly immune system that can easily eliminate attenuated in virulence bacteria at the wound site ^24^. Calculating the number of retrievable bacteria per fly we found that 29 of the 34 prototrophs (**Table 1**) and 17 of the remaining 44 strains (**Table 2 - Group A, C**) were able to colonize the flies like the wild type. In summary, we found 34 prototrophs (**Table 1**), 23 conditional prototrophs able to grow either in the host or in minimal medium cultures supplemented with fly extract (**Table 2, Group A-C**) and 21 auxotrophs unable to grow in the host or in culture (**Table 2, Group D**). Accordingly, 11.8% (57/482) of the core metabolic and 8.5% (8/94) of the non-metabolic attenuated strains were categorized as virulence-related (**Figure 5 A**), suggesting that many and functionally disparate metabolism genes are connected to *P. aeruginosa* virulence.

### Virulence-related metabolic genes of *P. aeruginosa* PA14 are part of core metabolism and are necessary for full virulence in mice

According to the KEGG database, the selected 57 virulence-related core metabolic genes belong to 7 metabolic categories and various metabolic gene pathways (**Figure 5 B**). Assessing single mutant representatives from each core metabolic pathway as defined in the KEGG database in a mouse lung infection model we found 15 mutants to be defective in virulence. The mutants belong primarily to amino acid metabolism [*PA14_57770 (hisC1), PA14_57780 (hisD), PA14_65250 (hisX), PA14_32690 (gtdA), PA14_16070 (hom)*], nucleotide metabolism [*PA14_17450 (surE), PA14_64220 (purD), PA14_52050 (purN)*] and the metabolism of co-factors and vitamins [*PA14_05590 (metF), PA14_07170 (epd), PA14_07600 (folk)*] (**Figure 6** and **Table 3**)

**Figure 6.**
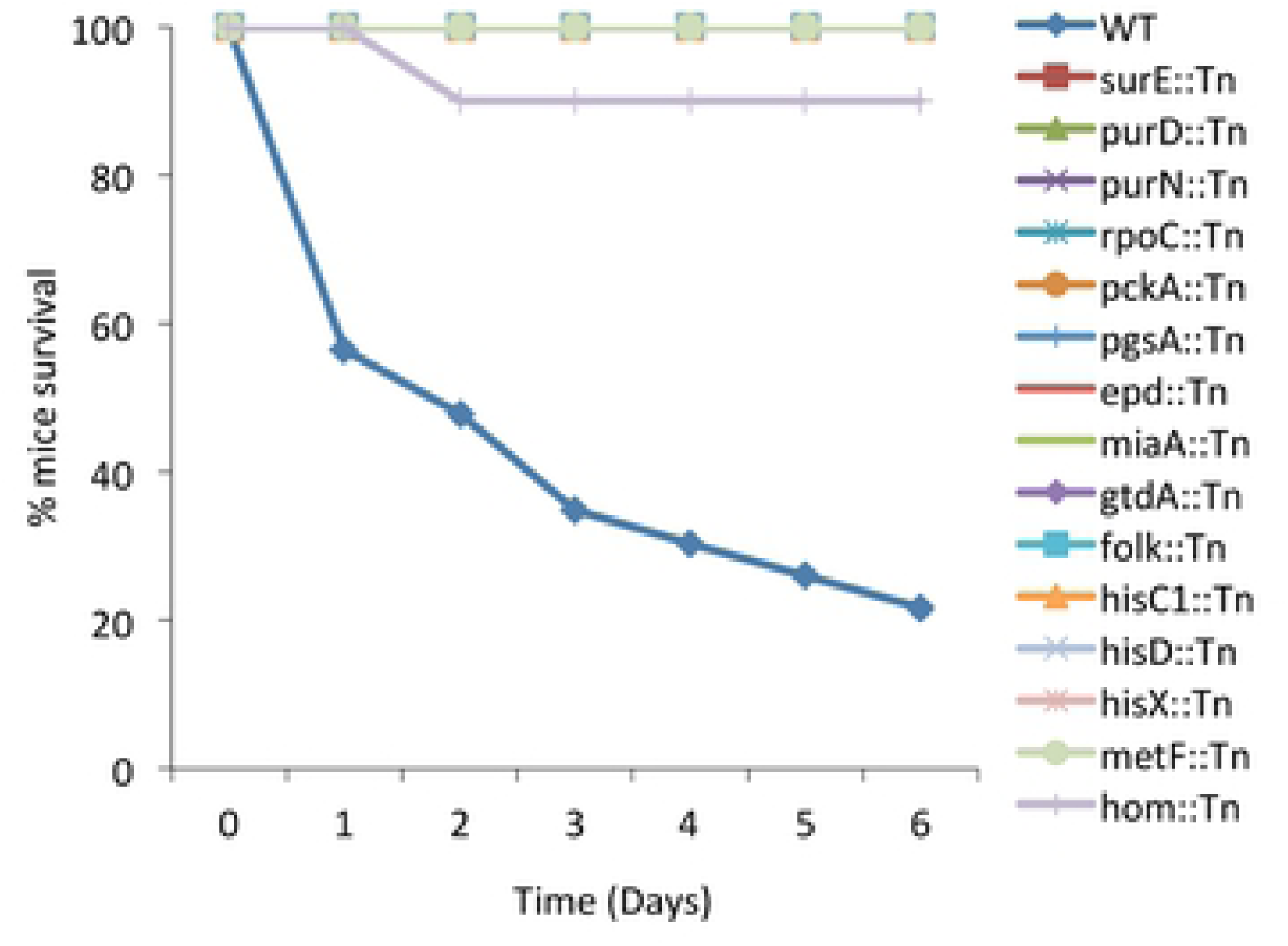
Survival kinetics of mice after intranasal inoculation with PA14 metabolic mutants. The graph shows the % mice survival after intranasal inoculation with 15 PA14 metabolic mutants and the wild type strain. Twenty-microliter aliquots of bacterial solution containing 2×10^7^ bacteria was administered intranasally to each mouse. Infected mice were monitored for 6 days (n=10-11).

**Table 3.** Pathways of the 15 PA14 metabolic genes corresponding to mutants found attenuated in mice lung infection. The names of the metabolic genes are listed in the first column. The second column shows the metabolic pathways in which each gene is involved based on the KEGG database, while the last column shows the major central metabolic pathways in which those pathways belong.

| # | Genes/Names | Major Metabolic Pathways<br>(based on KEGG database) | Major Central Metabolic Pathways |
| --- | --- | --- | --- |
| 1 | PA14_17450 (*surE*) | Purine, Pyrimidine & Nicotinate and<br>nicotinamide metabolism | Nucleotide metabolism<br>Metabolism of cofactors and<br>vitamins |
| 2 | PA14_64220 (*purD*) | Purine metabolism |  |
| 3 | PA14_52050 (*purN*) | Purine metabolism, One carbon pool by<br>folate |  |
| 4 | PA14_08780 (*rpoC*) | RNA polymerase<br>Protein families: genetic information<br>processing (Transcription machinery) |  |
| 5 | PA14_57770 (*hisC1*) | Histidine, Tyrosine & Phenylalanine<br>metabolism<br>Phenylalanine, tyrosine and tryptophan<br>biosynthesis etc. | Amino acid metabolism |
| 6 | PA14_57780 (*hisD*) | Histidine metabolism |  |
| 7 | PA14_65250 (*hisX*) | Histidine metabolism |  |
| 8 | PA14_32690 (*gtdA*) | Tyrosine metabolism |  |
| 9 | PA14_16070 (*hom*) | Glycine, serine and threonine<br>metabolism<br>Cysteine and methionine metabolism<br>Lysine biosynthesis |  |
| 10 | PA14_05590 (*metF*) | One carbon pool by folate | Metabolism of cofactors and<br>vitamins |
| 11 | PA14_07170 (*epd*) | Vitamin B6 metabolism |  |
| 12 | PA14_07600 (*folk*) | Folate Biosynthesis |  |
| 13 | PA14_68580 (*pckA*) | Glycolysis / Gluconeogenesis, Citrate<br>cycle (TCA cycle), Pyruvate metabolism | Carbohydrate metabolism |
| 14 | PA14_65320 (*miaA*) | tRNA dimethylallyltransferase<br>Protein families: genetic information<br>processing | Transfer RNA biogenesis |
| 15 | PA14_31700 (*pgsA*) | Glycerophospholipid metabolism | Lipid metabolism |

### Virulence-related metabolic genes of *P. aeruginosa* PA14 are compromised in various aspects of virulence

To better understand the connection of the multi-host attenuated metabolic mutants with virulence, we studied 13 of them for the production of virulence factors associated with acute infection, namely, bacterial motility, and T3SS and quorum sensing gene expression. Bacterial motility is very important for *P. aeruginosa* virulence ^64^. We examined the disruption of *P. aeruginosa* metabolic genes affecting its motility. We tested two types of motility: swarming and twitching for the 13 PA14 metabolic transposon (Tn) mutants found attenuated in both flies and mice. Mutants for *epd*, *surE*, *gtdA*, *purN*, *folK*, *metF*, *hom* and *rpoC*, were noticeably defective for swarming motility compared to the wild type strain (*P* < 0.001 for the first 7 and *P* < 0.01 for the last one), while *pgsA* and *purD*, exhibited increased swarming motility compared to the wild type (*P* < 0.05) (**Figure 7 A**). Regarding the twitching motility, the mutants *surE*, *gtdA*, *miaA*, *rpoC, pgsA* and *purN* exhibited a strong decrease in their ability to twitch compared to the wild type (*P* < 0.01) (**Figure 7 B**), but also the *pckA*, *folK* and *epd* mutants were significantly defective in twitching motility (*P* < 0.05) (**Figure 7 B**). Only *hisD* and *purD* Tn-mutants were not impaired in any of the two motilities.

**Figure 7.**
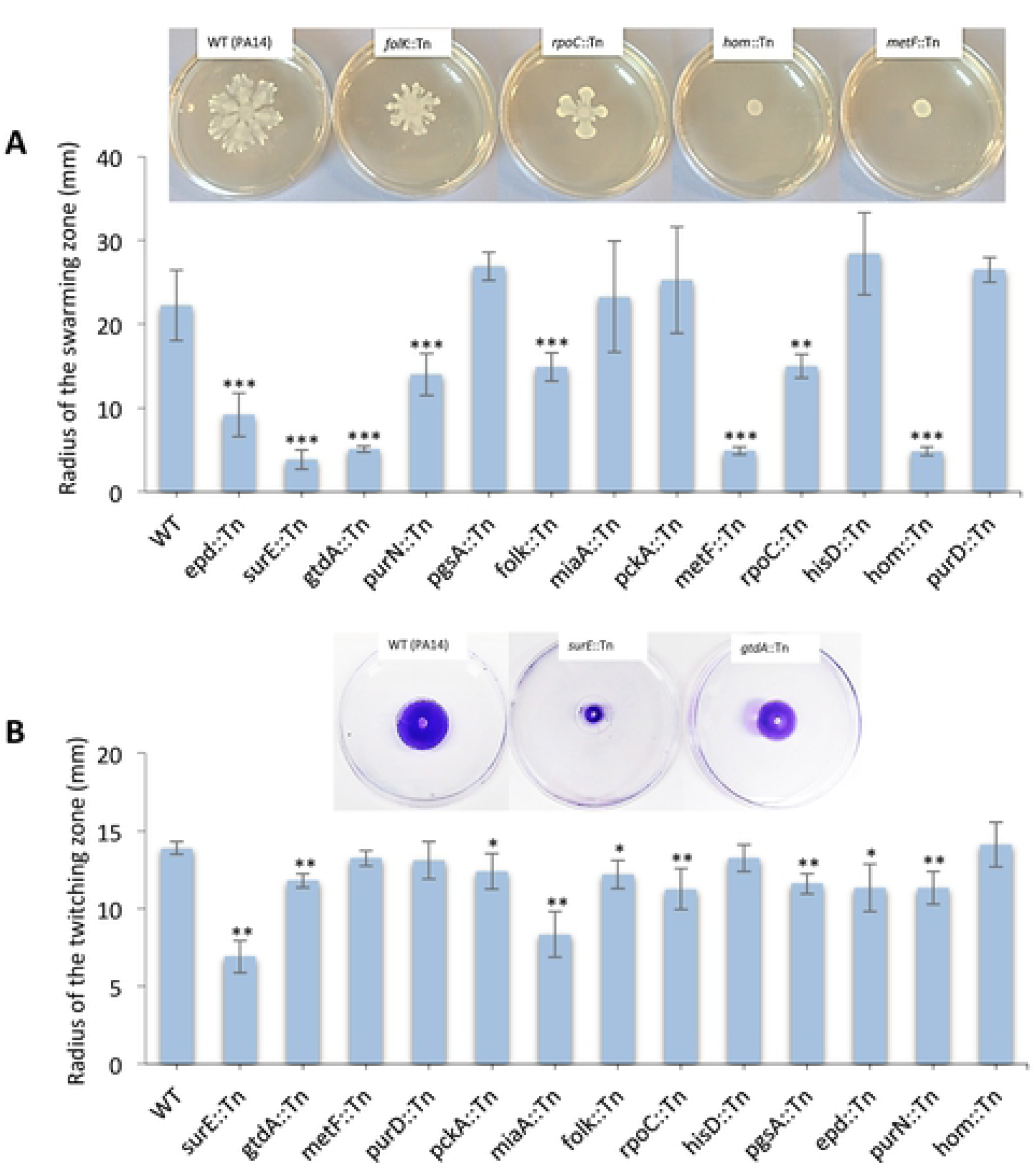
Swarming and twitching phenotypes of 13 PA14 metabolic gene mutants. (**A)** Swarming motility was measured by the length of the swarming zone from the center of the plate after inoculation of the cells at the center of the agar media and incubation at 37°C for 24 h. The results are from 3 independent experiments. n=3-13; **, P < 0.01; ***, P < 0.001 compared to WT by Mann-Whitney *U* test. Representative photos of swarm-negative phenotypes or defective swarming motility, of PA14 metabolic mutants compared to the wild type, are shown above the graph. (**B)** For the twitching motility the cells were stab-inoculated onto LB twitching plates (1% agar). The plates were incubated at 37 °C for 48 hours. The agar was removed, and the twitching zone was revealed by staining with crystal violet, measuring the maximum length of the twitching zone from the center of the plate. The results are from 3 independent experiments. n=4-7; *, P < 0.05; **, P < 0.01 compared to WT by Mann-Whitney *U* test. Representative photos of defective twitching motility, of PA14 metabolic mutants compared to the wild type, are shown above the graph. In both (A) and (B) error bars depict standard deviation.

Gram-negative bacterial pathogens have evolved multiple protein secretion systems that facilitate the infection of eukaryotic hosts. *P. aeruginosa* utilizes its type III secretion system (T3SS) to enhance its pathogenicity by injecting cytotoxic effector proteins into the host cells ^65^. T3SS expression is regulated transcriptionally and post-transcriptionally in response to host cell contact and environmental Ca^2+^ levels ^66^. We examined the expression of the T3SS regulatory gene *exsC* and the effector protein *exoT*, in the 13 Tn-mutants, under Ca^2+^ limiting conditions (5mM EGTA). We observed reduced expression of the *exoT* in 10 of the 13 selected mutants and overexpression in 1 of them, compared to the wild type (**Figure 8 A**). Similarly, the expression of *exsC* is reduced in 5 of these mutants, with only *purD* remaining unaffected in the expression of any of the two genes (**Figure 8 B**). Nevertheless, *purD* as well as the *miaA*, *rpoC*, *surE*, *purN* and *gtdA* mutants were compromised in the expression of the type IV pilus biogenesis gene *pilA* (**Figure 8 D**), while the *pa1L* gene, which is controlled by quorum sensing is significantly affected by *miaA, rpoC* and tentatively *purD* (**Figure 8 C**). *miaA*, is a tRNA isopentenyltransferase, a tRNA modification enzyme important for translation efficiency, while *rpoC* is the DNA-directed RNA polymerase beta subunit that has an important role in transcription. Our results suggest that the disruption in *miaA* and *rpoC* and the rest of the metabolic genes impact acute virulence by compromising one of more of key aspects of *P. aeruginosa* virulence.

**Figure 8.**
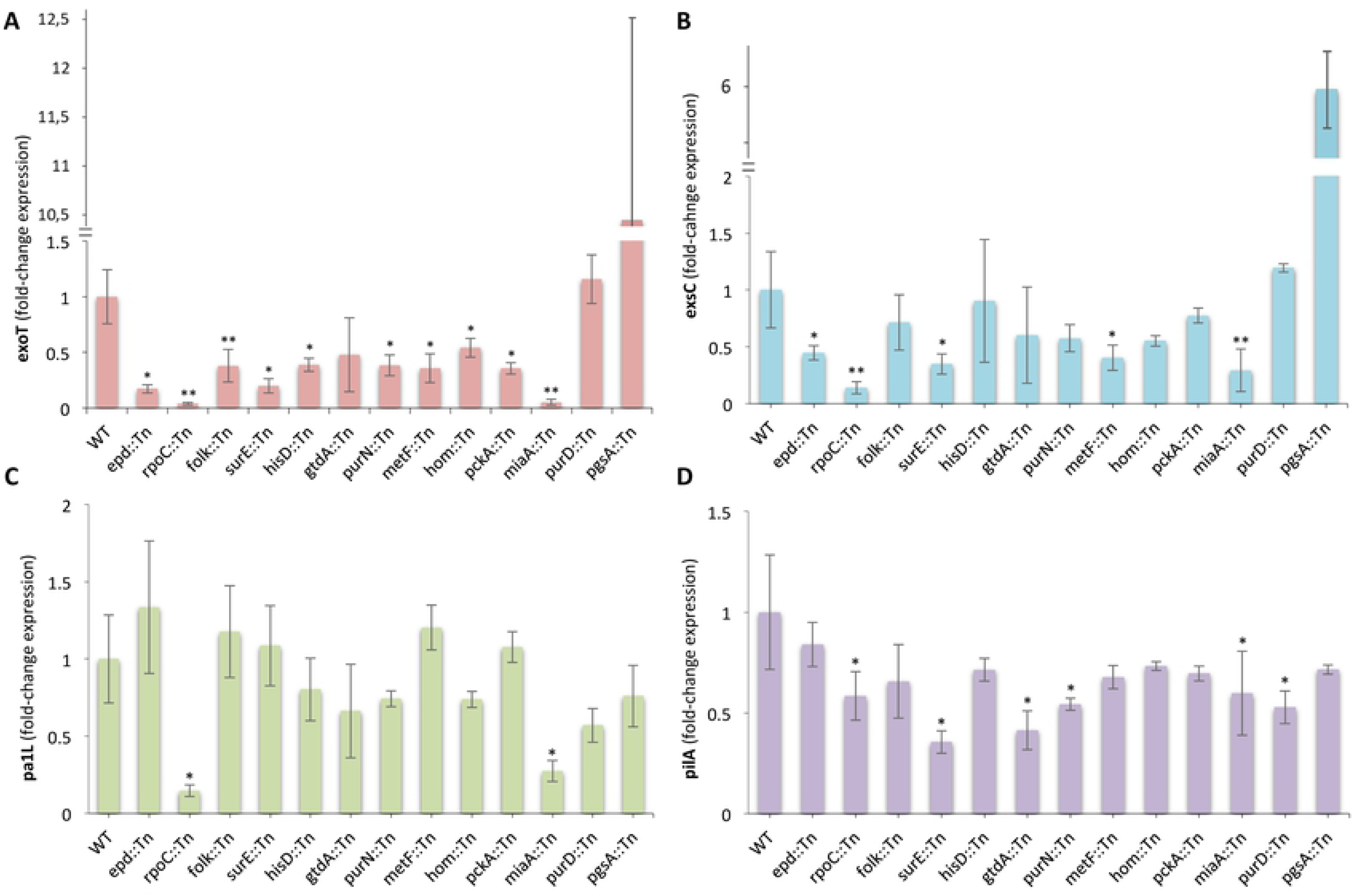
Expression of various virulence factors of 13 PA14 metabolic gene mutants. (**A, B**) Expression of T3SS genes in PA14 metabolic mutants. **(C, D)** Expression of *pa1L* and type IV pilus gene *pilA* in PA14 metabolic mutants. Bacteria were grown to an OD_600_ of 1 in LB with 5 mM EGTA before RNA extraction and cDNA synthesis. *, P < 0.05; **, P < 0.01 compared to WT by Mann-Whitney *U* test (n=3-7). In all plots (A-D) the average fold change ± standard deviation is shown.

### Virulence-related and “high” vs. “low” differentially expressed core metabolism genes are found in the same metabolic pathways

To pinpoint metabolic gene expression patterns important for *P. aeruginosa* virulence, we assessed the differential expression of the 78 core metabolism genes required for the full virulence of *P. aeruginosa* strain PA14 in the 3 selected high vs. the 3 selected low in virulence *P. aeruginosa* strains. Close inspection of differential gene expression in all possible “high” vs. “low” strain comparisons reveals that there is no consistent differential gene expression at any of the bacterial growth phases. That is, 45 of the 78 functionally important metabolic genes do not exhibit any differential expression among any of the 18 comparisons performed. The remaining 33, for which statistically significant up- or down-regulation was observed in at least one tested case, are mostly differentially expressed in the minority of “high” vs. “low” virulence comparisons (**Suppl. Figure 3**).

However, using the BioCyc Pathway/Genome Database Collection we identified 107 differentially expressed metabolic pathways containing genes differentially expressed between at least one highly (PA14, MTB-1, B136-33) vs. all 3 lowly (C3719, CF5, PACS2) virulent strains and pathways containing one or more of the 78 functionally validated genes. Overlapping these pathways, we find 8 of them containing genes upregulated in the all virulent strains against all low in virulence strains and at least one gene that compromises virulence when mutated (**Figure 9 A, Suppl. Table 6**). These common pathways are related to: (i) the 4-hydroxyl-phenylacetate degradation and succinate production, (ii) the glutamine biosynthesis from glutamic acid, (iii) the shikimate and chorismate biosynthesis from D-erythrose 4-phosphate, (iv) the branched chain amino acid biosynthesis of leucine, (v) the 2,5- and 3,5-xylenole degradation to citramalate, and (vi) the beta-oxidation of fatty acids (**Figure 9 B, Suppl. Table 6**). These data strongly support the idea that differences in virulence among *P. aeruginosa* strains, are arising from differential gene expression of genes belonging in specific core metabolism pathways.

**Figure 9.**
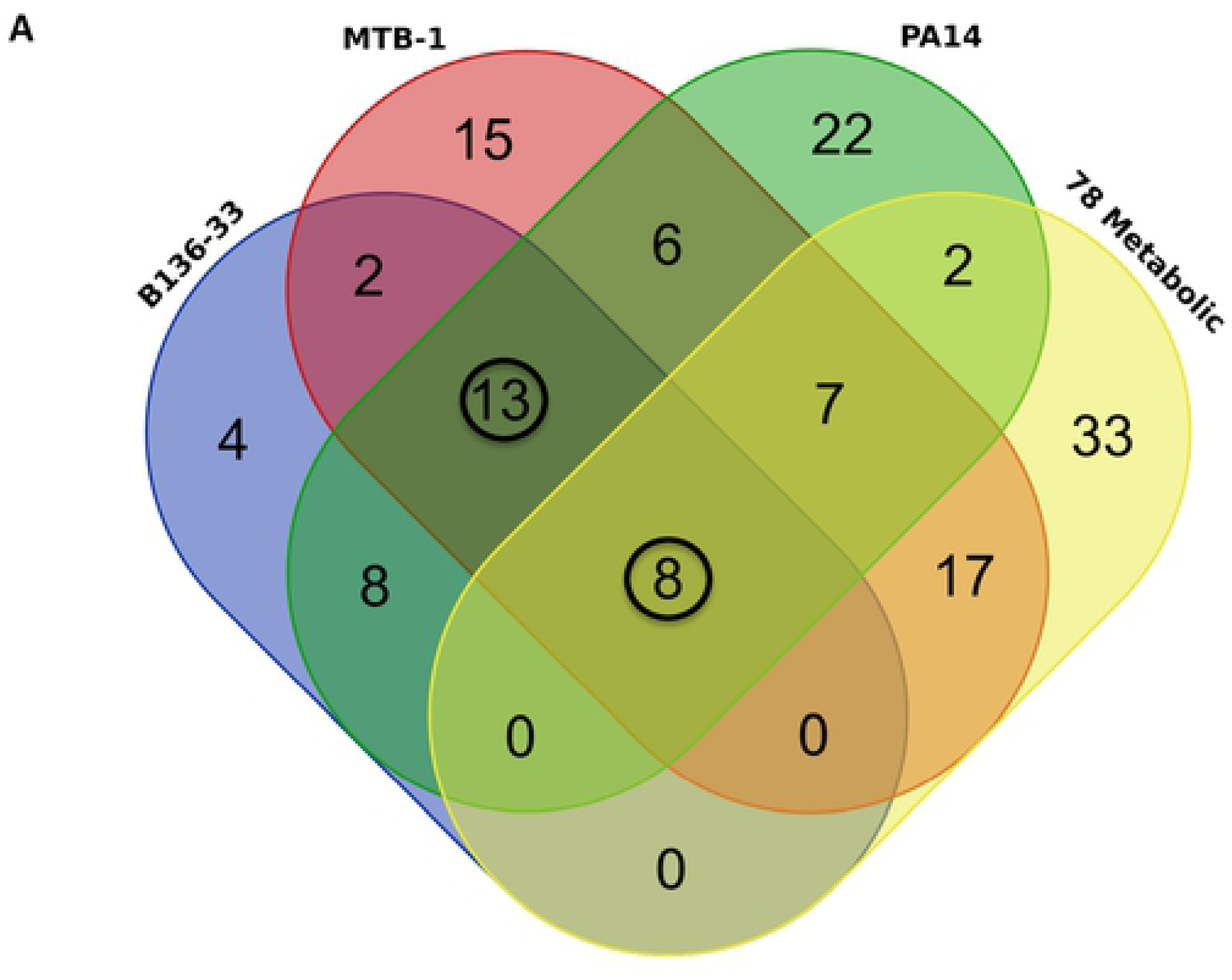
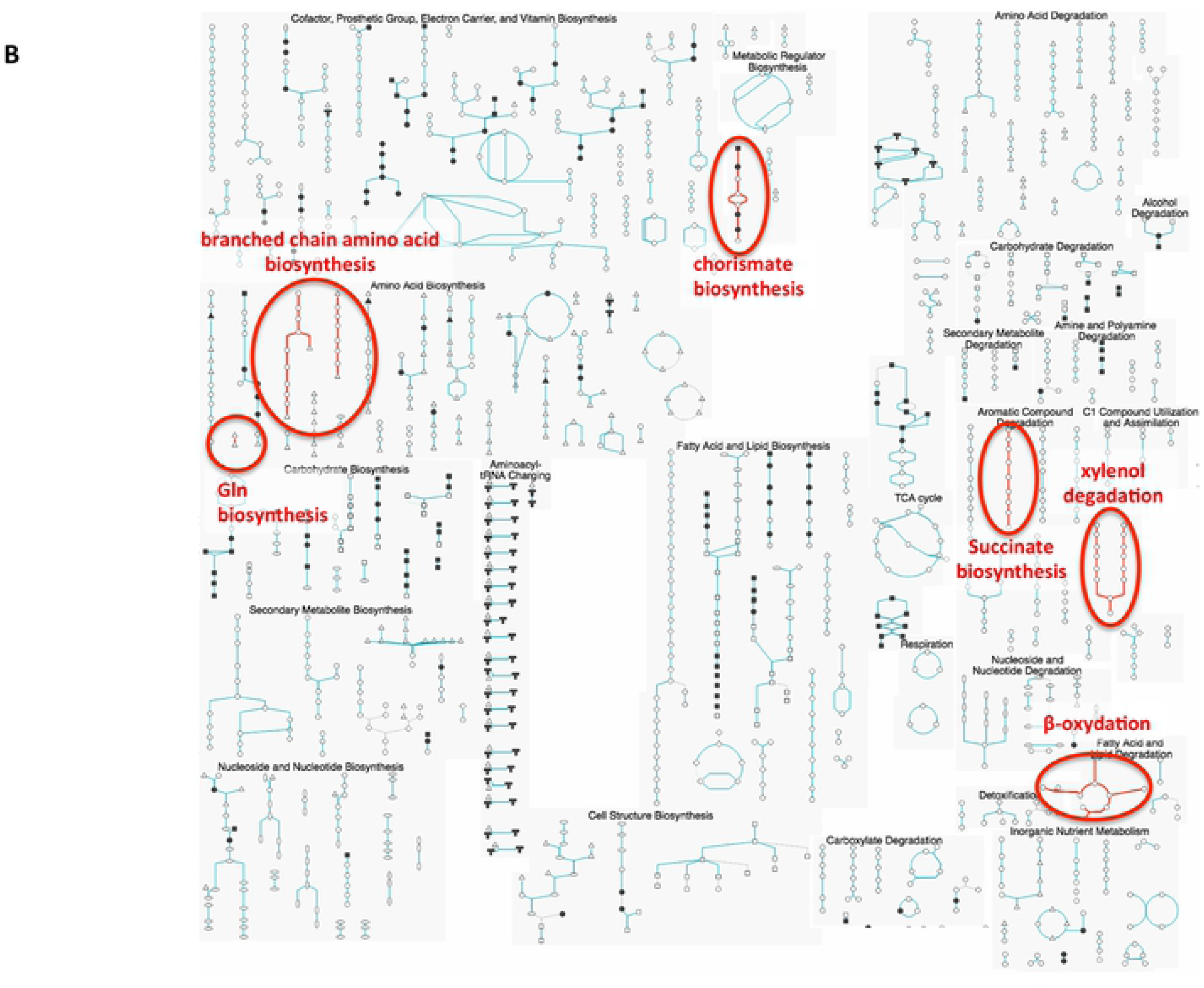
(A) Overlap among 4 metabolic gene pathway groups. Three expression-based metabolic pathway groups arise from differentially expressed genes comparing each of the 3 high (B136-33, MTB-1 or PA14) with all 3 low (CF5, C3719 and PCS2) in virulence strains, while the 4th functionality-based metabolic pathway group arises from the 78 genes functionally important for full virulence in flies. Out of totally 137 metabolic pathways assessed for overlap based on BioCyc Database Collection (https://biocyc.org) 21 were common to the 3 expression-based metabolic pathway groups, while 8 of them were also functionally important for virulence. We circle the numbers of the 8 functionality-based plus 13 more, totaling 21 common expression-based metabolic pathways. **(B) Common metabolic pathways implicated in *P. aeruginosa* virulence.** Hihglighted are the common pathways shown in (A) within the Cellular Overview of *P. Aeruginosa* PA14 metabolism from BioCyc.

## Discussion

Aspects of bacterial metabolism have been linked to virulence, including the T3SS ^67^, the regulation of exotoxin A expression ^68,69^ and the adaptation to the host nutritional environment ^70^. Carbon source availability, for example, affects virulence genes presumably as an adjustment to the host environment^71^. *P. aeruginosa* gene AaaA is an arginine-specific autotransporter providing a fitness advantage in chronic wound infections where the sole source of nitrogen is peptides with an aminoterminal arginine ^72^. In addition, *P. aeruginosa* CbrA is a sensor kinase involved in carbon and nitrogen utilization, which affects swarming, biofilm formation, cytotoxicity, and antibiotic resistance ^73^. Computational screening further elucidates interconnectivity between *P. aeruginosa* virulence factor synthesis and growth with a metabolic network as a denominator ^18^. Such studies support the notion that core metabolism genes are growth-essential and unsuitable drug targets due to their high potential for resistance development ^18^. Our functional transcriptomics approach though suggests the existence of a widespread growth-independent effect of core metabolism gene expression in virulence.

We defined virulence genes as those necessary to the full virulence of *P. aeruginosa* in conditions able to sustain their growth or their host colonization ability. Thus, we included as growth-independent, mutants able to grow similarly to wild type strain in minimal media supplemented or not with fly extract; and mutants able to colonize the host in the infection assay (wound or oral) supporting their atteniuation in virulence. t We found that 11.8% of the growth-independent metabolic genes and 8.5% of the non-metabolic genes are directly linked to virulence. Of the former, histidine metabolism were conditional prototrophs for being able to grow efficiently in minimal media supplemented with fly extract and colonize the flies as efficiently as the wild type *P. aeruginosa* strain. The virulence attenuation of these mutants is more likely due to defects in virulence rather than growth defects. Similarly, most of the purine metabolism mutants were conditional prototrophs for being able to grow in minimal media that contained fly extract, but these mutants were unable to colonize flies to the same extent as the wild type strain. This suggests that although enough nutrients for growth exist in the host tissues, bacteria are unable to acquire them through their virulence factors. Interestingly, we also identified conditional prototrophs that although unable to grow efficiently in culture in any of the two types of minimal media, they could colonize flies normally, thus the corresponding genes may help to damage the host, irrespective of their contribution to growth.

To validate our approach, we assessed VF production in a subset of the metabolic gene mutants that were attenuated in virulence not only in flies but also in a mouse lung infection model. We found that all of them exhibit defects in at least one aspect of virulence. Moreover, most of them are involved in metabolic steps absent from the human metabolism network. Thus, many metabolic genes could be targets for anti-infective therapies against acute *P. aeruginosa* virulence. It is nevertheless questionable whether prioritization in pharmacological targeting should be given to virulence-related metabolic genes also affecting growth or those that do not affect growth.

Moreover, we found differential expression of many known virulence factors and some metabolism genes through classical transcriptome analysis of highly vs. lowly pathogenic *P. aeruginosa* strains. Combinatorial functional transcriptomics analysis at the pathway level was much more informative, revealing metabolic pathways containing differentially expressed genes between all 3 highly virulent vs. all 3 lowly virulent strains and genes required for full virulence: (i) the 4-hydroxyl-phenylacetate degradation and succinate production, (ii) glutamine biosynthesis from glutamic acid, (iii) shikimate and chorismate biosynthesis from D-erythrose 4-phosphate, (iv) superpathway of branched chain amino acid biosynthesis of valine, leucine and isoleucine, (v) 2,5- and 3,5-xylenole degradation to citramalate, and (vi) beta-oxidation of fatty acids. Thus, *P. aeruginosa* virulence can be analysed at the trancriptome and functional level using common core metabolism modules that control and indicate the virulence of disparate *P. aeruginosa* strains.

While the impact of specific core metabolism genes is not discernible at the genomic level, interesting patterns emerge. By classifying the 30 fully sequenced *Pseudomonas* species experimentally in two different *Drosophila* infection assays (wound and oral infection) as low, medium or high in virulence, we found that the phylogeny of the 30 strains correlates well with their virulence classification in the wound infection, but not in the oral infection assay. This is probably because the wound infection is more acute and guided by highly efficient virulence related genes, as opposed to the intestinal environment in which *Pseudomonas* is less adapted. The phylogeny of the 30 strains also correlates well with virulence factor gene content, which is expected since *P. aeruginosa* species have their own repertoire and group together in the phylogenetic tree and separately from the non-*P. aeruginosa* strains. Similarly, the phylogeny of the 30 strains also correlates with major metabolic pathway gene content. Thus, the existing correlations between pathogenicity of the 30 *Pseudomonas* strains and gene content cannot be attributed specifically to VFs or metabolic genes. The gene content of the 30 *Pseudomonas* spp. regarding five metabolic pathways (carbon, glycolysis, pyrimidine, propanoate metabolism and valine, leucine, isoleucine degradation) explains their virulence classification upon the wound infection, but not the oral infection. Nevertheless, when the analysis is repeated with only the *P. aeruginosa* strains, no correlation is evident for any comparison. Thus, the existing correlation between the pathogenicity of the 30 *Pseudomonas* strains and genomic content in metabolic genes cannot be attributed to specific genes, but rather to broader interspecies differences. Given the difference in expression of virulence factors and core metabolism pathways in distinguishing highly from lowly virulent strains it is likely that gene promoter differences rather than gene content of VFs and core metabolism genes accounts for the strain to strain variation in virulence potential.

## Acknowledgments

Bacterial RNA sequencing was performed at the BSRC Al. Fleming Genomics Facility. We thank Dr Vaggelis Harokopos for the NGS experiments and Dr Martin Reczko for initial bioinformatic analyses. We also thank Stelios Theofilou and Eleni Ioannidou for helping with the *Drosophila* infection screens.

## Supplementary data

**Suppl. Figure 1.**
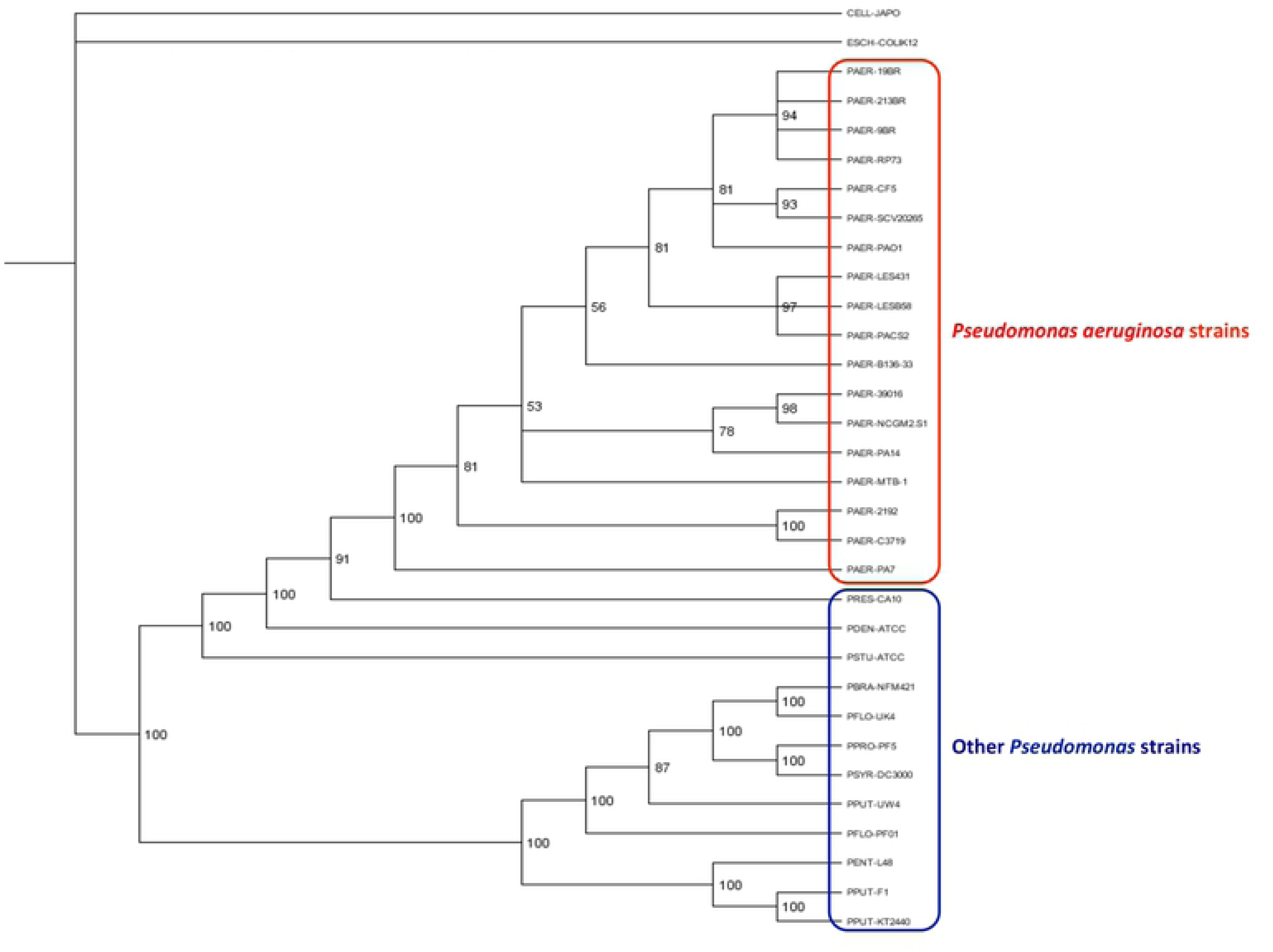
Phylogenetic tree including the 18 *P. aeruginosa* and the 12 other Pseudomonas strains clustering separately. The tree was constructed using the sequences of the conserved gene gyrB and the sequences of 16S rRNA and the Bayesian Model (MrBayes, http://brahms.biology.rochester.edu/software.html, 2001).

**Suppl. Figure 2.**
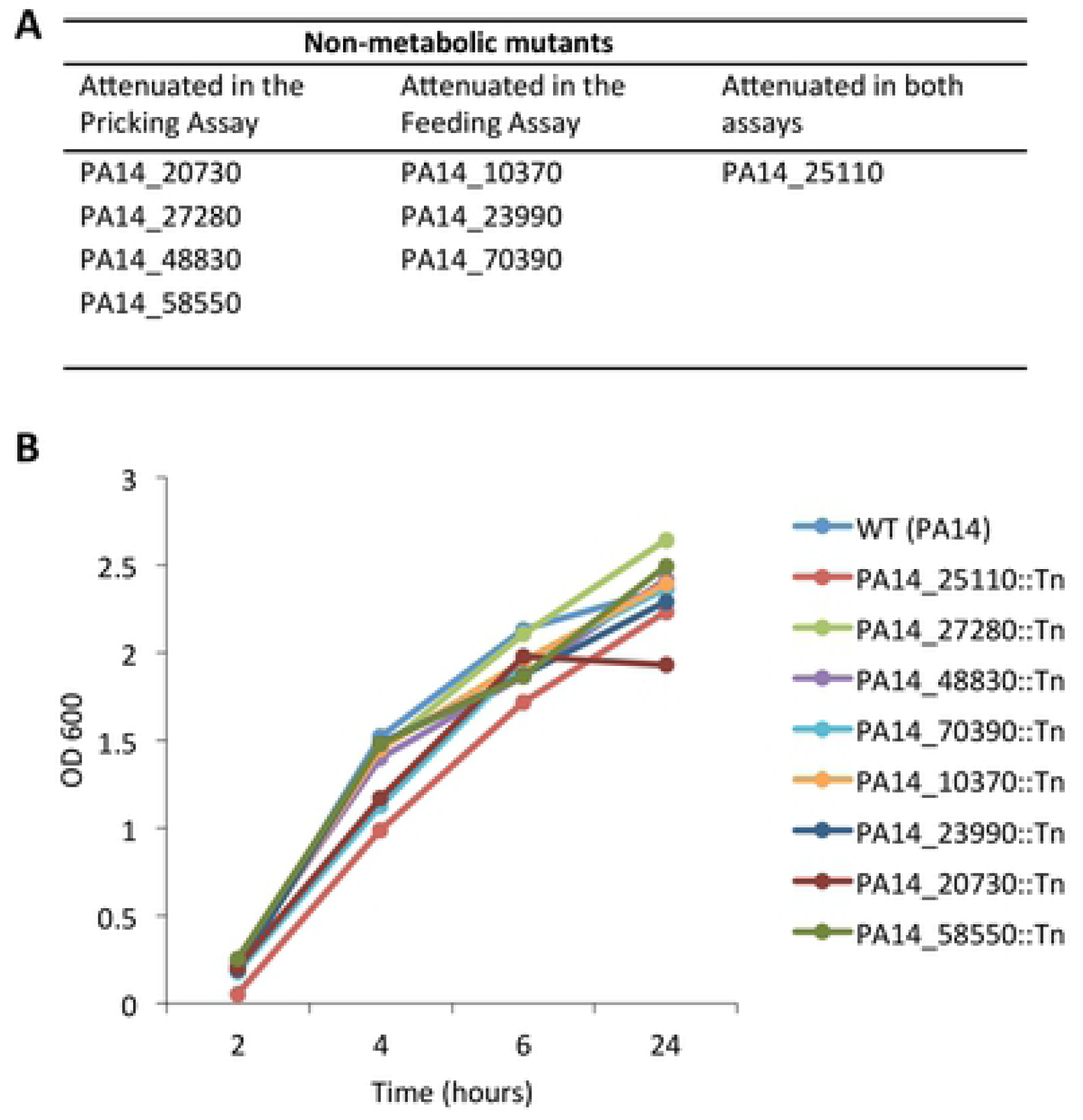
Selected non-metabolic PA14 mutants and assessment of their growth in glucose minimal media supplemented with S% fly extract. **(A)** Non-metabolic PA14 transposon mutants that found attenuated in flies during wound and/or oral infection. The table shows all the virulence-related non-metabolic mutants and the assay in which were found attenuated. **(B)** Growth of selected non-metabolic PA14 mutants in glucose minimal media supplemented with 5% fly extract. The growth of the selected non-metabolic mutants was assessed only in glucose minimal medium that additionally contained 5% fly extract to verify that these mutants have the potential to grow in flies. The optical density was measured at four time points. All of them were able to grow in this medium at the same extent as the wild-type PA14.

**Suppl. Figure 3.**
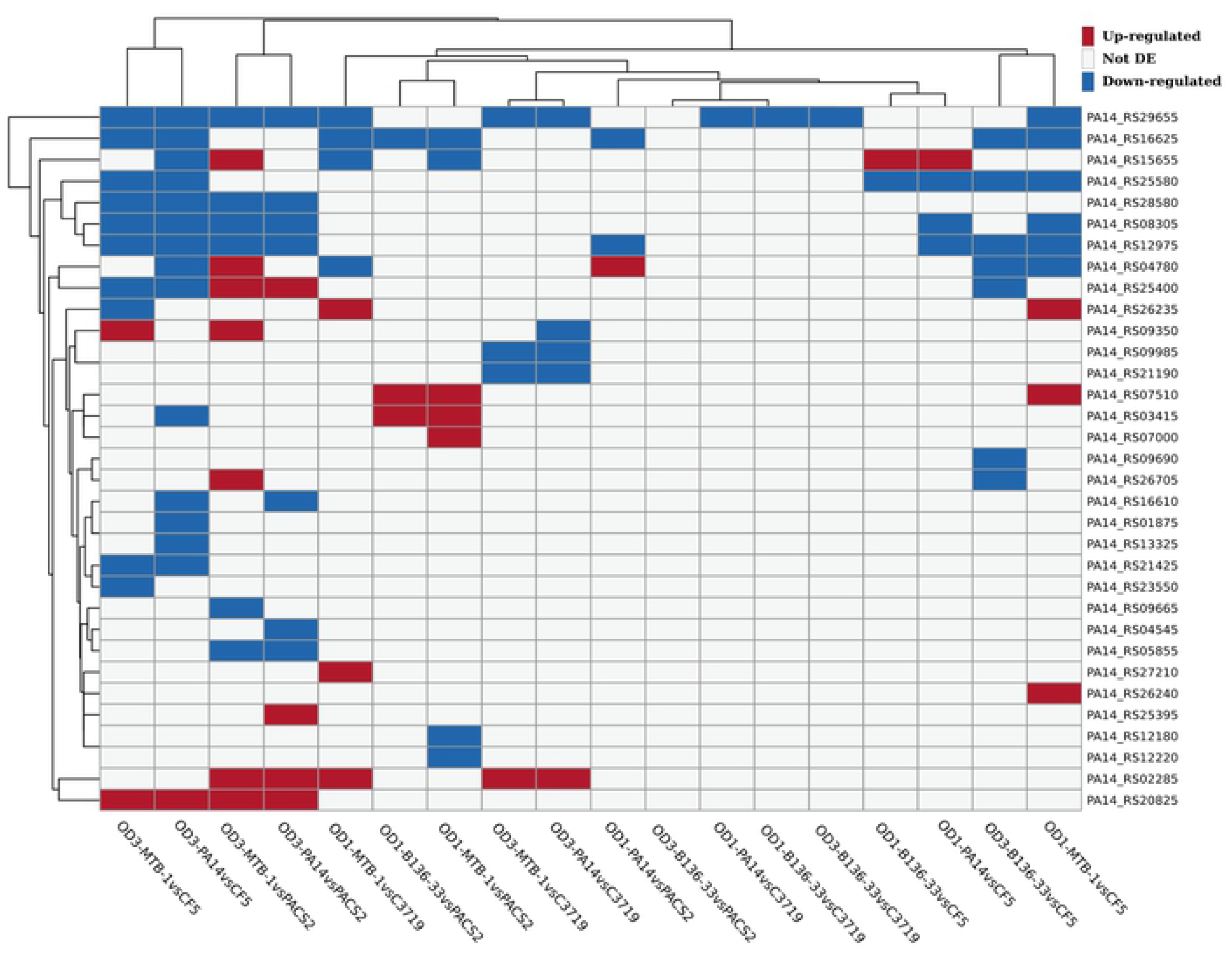
Differential gene expression of core metabolism genes compared between 18 different high vs. low in virulence conditions. Display of the 33 genes significantly up-(red boxes) or down-regulated (blue boxes) in at least one of the 18 comparisons performed. 26 of the 33 genes alter their expression in less than S comparisons. No centering or scaling was applied to rows and columns of the data matrix. Both rows and columns were clustered using Manhattan distance and average linkage hierarchical clustering.

**Suppl. Figure 4.**
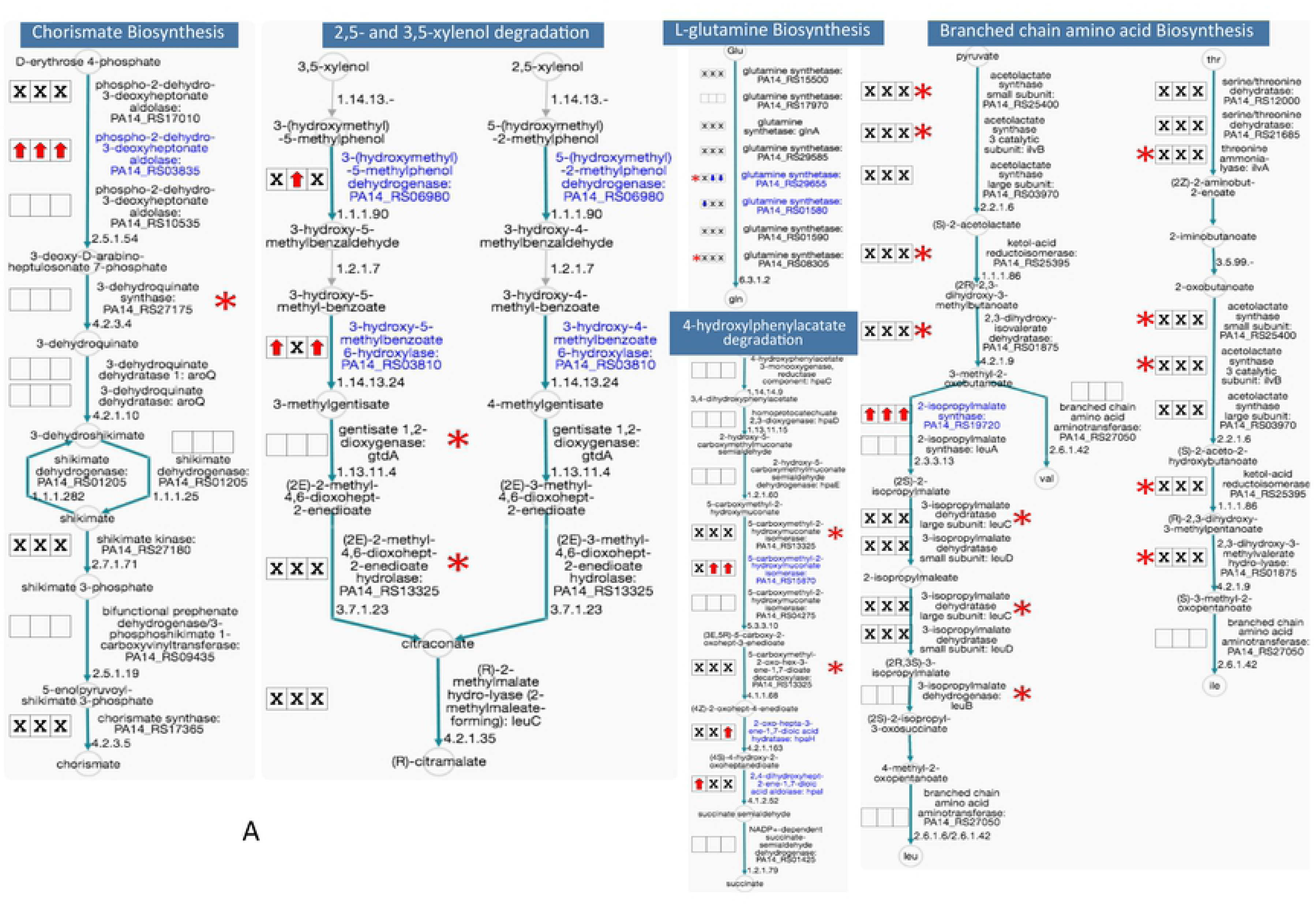
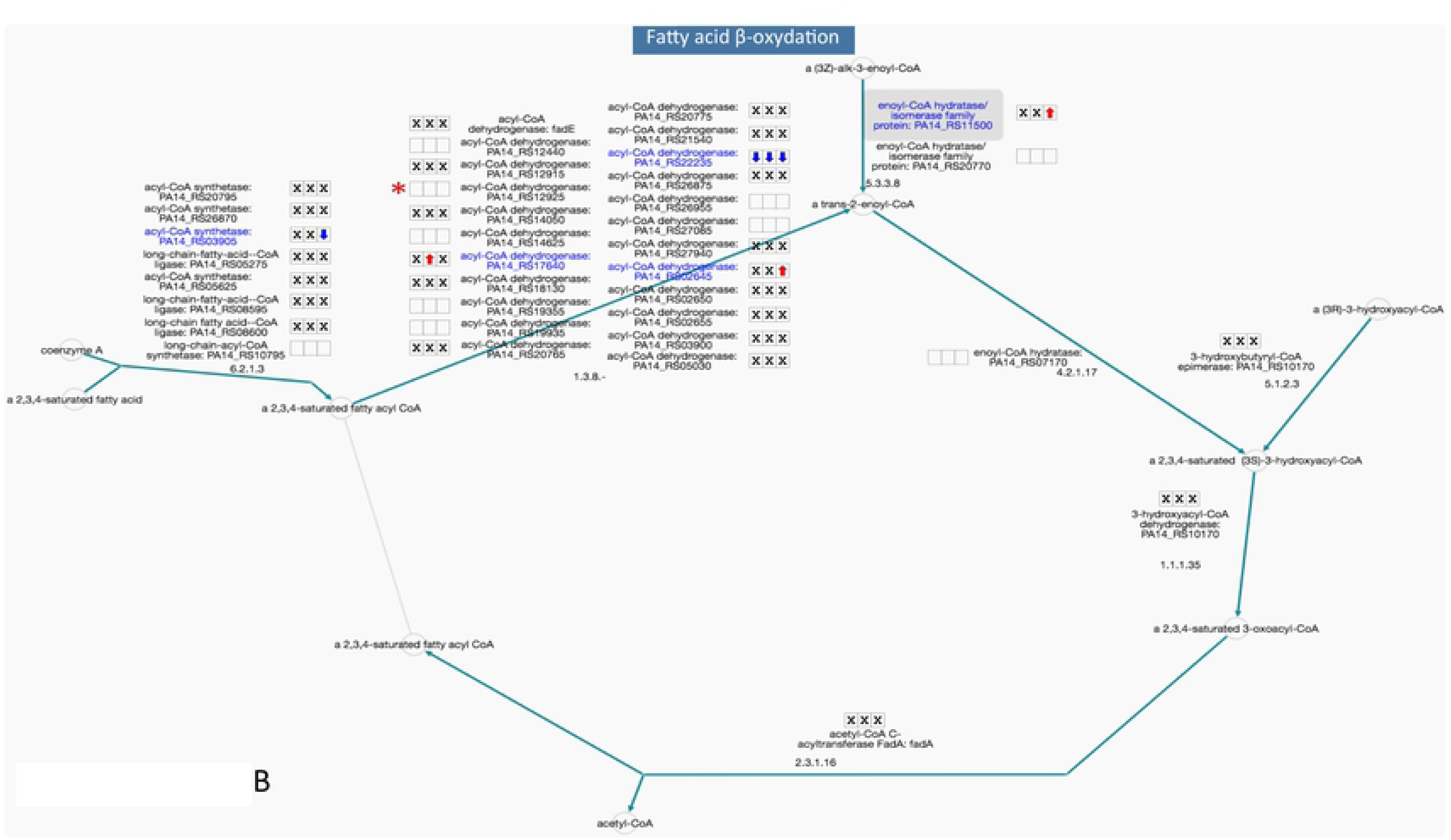
(**A,B**) Pathways with core metabolism genes with implications in virulence (red stars) and at least one gene (named in blue) differentially expressed in a consistent manner in comparisons of at least one highly virulent strain to all low virulent strains. All genes are labeled according to their differential expression patterns with boxed symbols: empty - no differential expression; red arrow - Upregulated; blue arrow -downregulated; X - conflicting differential expression. The 3 boxes correspond (left to r ight) to strains B136-33, MTB-1 and PA 14 respectively. The L-leucine biosynthesis pathway is depicted as part of the superpathway of branched chain amino acid biosynthesis.

